# Programmable downsizing of CRISPR-Cas9 activity for precise and safe genome editing

**DOI:** 10.1101/2020.10.31.361733

**Authors:** Masaki Kawamata, Hiroshi I. Suzuki, Ryota Kimura, Atsushi Suzuki

## Abstract

CRSIPR-Cas9 system has opened up the avenue to efficient genome editing^1–4^. However, together with known off-target effects, several concerns of current CRISPR-Cas9 platform, including severe DNA damage, cytotoxicity, and large genomic alteration, have emerged in recent reports^5–7^ and establish a formidable obstacle to precisely model allele dosage effects of disease mutations and risk variants, especially mono-allelic effects, and correct them. Here, by developing an allele-specific indel monitor system (AIMS), we demonstrate that small and simple modification of conventional single-guide RNAs (sgRNAs) enable programmable tuning of CRISPR-Cas9 activities and alleviate such adverse effects. AIMS, which visualizes various indel events in two alleles separately in living cells, is convenient and accurate to determine the *in vitro* editing efficiency and revealed frequent mosaicism during genome editing. Using AIMS, we show that adding cytosine stretches to the 5’ end of conventional sgRNA efficiently reduced Cas9 activity in a length dependent manner. By combining systematic experiments and computational modeling, we established the quantitative relationships between the length of cytosine extension and multiple aspects of CRISPR-Cas9 system. In general, short cytosine extension dramatically relieves p53-dependent cytotoxicity and suppression of homology-directed repair (HDR) while relatively maintaining on-target activity. Long cytosine extension further decreases on-target activity, thereby maximizing mono-allelic editing, while conventional system typically induces bi-allelic editing. Furthermore, such downregulation of on-target activity contributes to downregulation of relative off-target activity and protection of HDR-allele from second off-target editing. Therefore, cytosine extension method finally enables both single-step generation of heterozygous single-nucleotide disease mutations from homozygous states in mouse ES cells and correction of heterozygous disease mutations in human iPS cells. Taken together, our study proposes updates of standard CRISPR-Cas9 platform in mammalian cells toward precise and safe genome editing in diverse applications.

## Introduction

CRSIPR-Cas9 system has opened up the avenue to efficient genome editing^1–4^. It is still challenging to model and correct most genetic variants that contribute to various diseases. Several methods such as CORRECT combine the standard CRISPR-Cas9 system and homology-directed repair (HDR) for this purpose^8,9^. However, they typically require introduction of silent mutations in HDR templates to protect the HDR-allele from second editing since the HDR templates are easy targets for off-target activities of Cas9, and multiple cloning steps are thus inevitable to revert silent mutations. Other approaches such as base editing (BE) or prime editing (PE) avoid DNA double-strand break (DSB)^10–13^, but still accompany with insertions/deletions (indels) and undesirable mutations caused by editing errors^14–17^. Besides known off-target effects, multiple recent studies have unveiled that several adverse effects, such as cytotoxicity with severe DNA damage, large on-target genomic deletion, and chromosomal rearrangement, are prevalent in mammalian cells^5–7^. Indeed, the large deletions are induced in up to 20% of cells^7^. Among them, the large DNA deletions are particularly underestimated in the next-generation sequencing (NGS)-based methods, which are typically used for development of CRISPR-based technologies, such as CORRECT, BE, and PE, because short read sequencing analyzes only partially matched target sequences. NGS-based methods also miss clonal and allelic editing information, including the prevalence of mosaicism. To investigate an avenue for safe genome editing, we first attempted to establish a convenient but accurate experimental platform to visualize the dynamics of genome editing in each single allele at the single cell level in living cells: allele-specific indel monitor system (AIMS).

## Results

### Allele-specific indel monitor system (AIMS)

An allele-specific indel monitor system (AIMS) employs insertion of the monitor cassette containing 2A self-cleaving peptides (P2A) and two distinct fluorescent proteins (tdTomato and Venus) in two alleles (Fig. 1a). Use of sgRNA targeting a P2A sequence allows us to analyze indel induction in a pair of alleles by two distinct colors in real-time and at clonal level without sequence analysis. By inserting AIMS cassette downstream of coding regions of genes, which products localize to the nuclei or cell membrane, e.g. transcription factors (TF) or membrane proteins (MP), changes in localization of two fluorescence can distinguish nine combinations consisting of in-frame indels, frameshift indels or large deletions, and no indel at each allele (Fig. 1a). For example, in-frame indels disrupt endopeptidase recognition of P2A peptides and thus result in change in fluorescence localization to nucleus or membrane. AIMS is also sensitive to large deletions, which cause loss of fluorescence. In AIMS, multiple sgRNAs can be tested using several target sites in P2A sequence and/or generating P2A variants with silent mutations (Fig. 1b). In this study, we used an original P2A sequence, named P2A_1_^18^, and one of its variants, named P2A_2_, and tested six sgRNAs targeting P2A_1_ or P2A_2_ (Fig. 1b). We first developed AIMS in mouse embryonic stem cells (mESCs) by targeting T-Box transcription factor 3 (*Tbx3*) and membrane protein E-cadherin (*Cdh1*) because they are homogenously expressed in mESCs under a 2iL culture condition^19,20^ (Extended Data Fig. 1a). AIMS successfully distinguished various combinations and was consistent with sequence validation (Fig. 1c and Extended Data Fig. 1b, c).

**Fig. 1.**
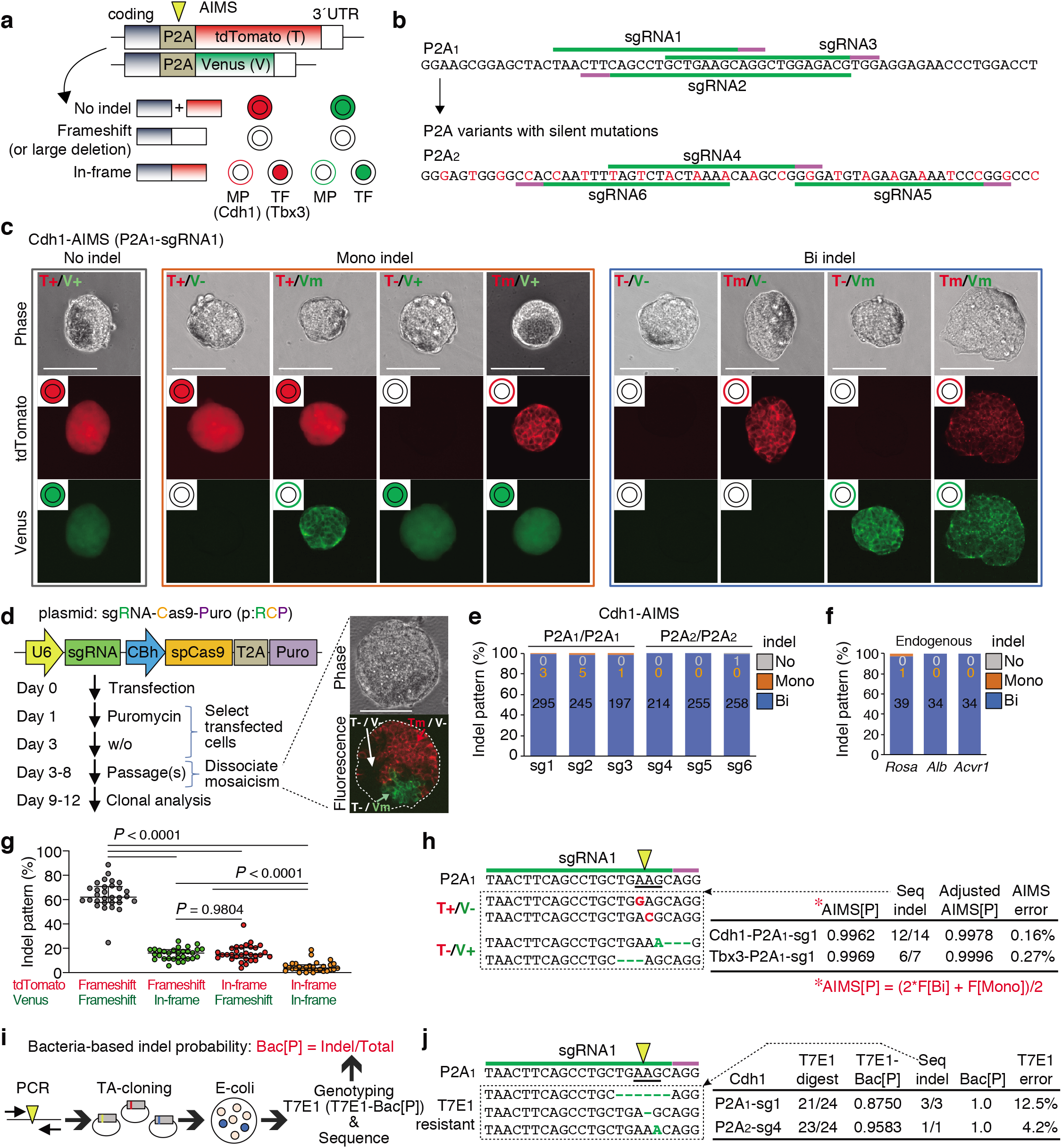
Visualization of allele-specific genome editing events by AIMS. **a**, Schematic of AIMS. A mESC clone harboring dual color reporters is generated by fusing P2A-fluorescence cassettes at the ends of coding regions of target genes. P2A is targeted by sgRNA-Cas9 (yellow arrowhead). TF, Transcription factor; MP, Membrane protein. **b**, Target sequences of sgRNAs in the P2A1 and P2A2 are shown. The original P2A is denoted as P2A1, and the variant generated by silent mutations (red) is denoted as P2A2. **c**, Representative results of Cdh1-P2A1-AIMS. Genotypes are determined by nine combinations of tdTomato/Venus expression and localization. T, tdTomato; V, Venus; +, no indel; m, in-frame indel represented by membrane localization; -, frameshift indel or large deletion represented by loss of fluorescence. Scale bar indicates 100 μm. **d**, Schematic description of protocol for genome editing using all-in-one CRISPR plasmids (pRCP, sgRNA-Cas9-Puro). The images show mosaicism in a single cell-derived primary puromycin-resistant colony. Scale bar indicates 100 μm. **e**, Indel patterns are measured by Cdh1-P2A1-AIMS and Cdh1-P2A2-AIMS. sg1-6, sgRNAs1-6 shown in Fig. 1b. Data are shown as mean from n = 3 independent experiments performed at different times, except for sg1 (*n* = 6). Total number of clones analyzed is shown in each column (also in **f**). **f**, Indel patterns for the endogenous genes are determined by sequence analysis at the clonal level. Data are shown as mean from *n* = 3 independent experiments performed at different times. **g**, Percentages of the four types of bi-allelic indel patterns are shown. Dots indicate individual data points (*n* = 30, 6 sgRNAs, Tbx3- and Cdh1-AIMS) and median with interquartile range are shown. Statistical significance is assessed using Welch’s test with post hoc Games–Howell test. **h**, Representative indel sequences in the P2A1 region of tdTomato or Venus allele in the T+/V- or T-/V+ clones, respectively (left) and AIMS error rates (right) are shown. AIMS[P], the value of indel probability calculated from frequency (F) of bi-allelic [Bi] and mono-allelic [Mono] indel clones. The formula is shown in red below. Seq-indel, the exact number of bi-allelic indel clones with a phenotype of T+/V- or T-/V+ is determined by sequencing the tdTomato or Venus allele, respectively. Adjusted AIMS[P], indel probability considering the Seq-indel data. AIMS error, the values of Adjusted AIMS[P] minus AIMS[P]. **i**, Schematic description of the procedure for calculating bacteria-based indel probabilities (Bac[P]). **j**, Representative T7E1-insensitive indel sequences (left) and the error rate of the T7E1 assay (right) are shown. T7E1-Bac[P], indel probability calculated from the rate of clone number sensitive to T7E1 digestion. Seq-indel, the exact number of indel clones is determined by sequencing the PCR products which are not digested by T7E1. Bac[P], indel probability considering the Seq-indel data. T7E1 error, the values of Bac[P] minus T7E1-Bac[P]. Arrowheads and underlines indicate the position of the DSB sites and codon, respectively (**h, j**).

To enhance experimental reproducibility, we mainly utilized the all-in-one plasmids expressing single-guide RNA (sgRNA), Cas9, and puromycin-resistant cassette (p:RCP), and performed AIMS analysis in cells selected by puromycin treatment (Fig. 1d). Notably, approximately 30 % of primary colonies derived from puromycin-resistant single cells exhibited mosaicism (Fig. 1d). Thus, the primary colonies were dissociated and the secondary ones with homogenous fluorescent pattern were analyzed (Fig. 1d). Bi-allelic indels were induced in more than 99.4% of mESC clones for all of six tested sgRNAs targeting P2A_1_ or P2A_2_ (Fig. 1e). Similar results were obtained when endogenous genes were targeted (Fig. 1f). Although the CRISPR editing efficiency is reportedly influenced by chromatin accessibility^21^, we confirmed high rates of bi-allelic indels for *Alb* gene, not expressed in mESCs.

Next, we investigated the accuracy of AIMS. When analyzing bi-allelic indel clones, allelic bias in both indel induction and frameshift/in-frame indel frequency was not evident (Fig. 1g and Extended Data Fig. 1d). Frequencies of in-frame indels were lower than those of frameshift indels or large deletions. Next, the error rates of AIMS data-based indel probability (AIMS[P]) were investigated by additional sequence analysis of the rare population of the tdTomato^+^/Venus^indel^ and tdTomato^indel^/Venus^+^ heterozygous clones (Fig. 1h). The 86% of these ostensibly heterozygous clones were turned to be homozygous, resulting in the error frequency less than 0.3% (Fig. 1h). We next performed a standard T7E1 survey assay with a bacterial cloning process in our experiments, determined indel probability (T7E1-Bac[P]), and estimated the error rates (Fig.1i). Importantly, additional sequence analysis revealed that approximately 8% of indels were not digested by T7E1 (Fig.1j), suggesting that AIMS is more accurate than T7E1 assay. Thus, combination of T7E1 and sequence analysis was performed hereafter when determining bacterial cloning-based indel probability (Bac[P]). These results collectively suggest that the current CRISPR-Cas9 system basically induces bi-allelic DNA cleavage when appropriate sgRNAs are designed and Cas9-introduced cells are sufficiently selected, and that editing outcomes at each allele are stochastic and highly dynamic, leading to frequent mosaicism.

### Downsizing of sgRNA-Cas9 activity by sgRNA modification

Consistent with our findings, to generate heterozygous genotypes, other methods such as CORRECT have basically employed bi-allelic editing but introduced some technical approaches, such as the use of mixed HDR templates, to control editing outcomes of two alleles for heterozygosity^8,9,22^. We alternatively attempt to maximize mono-allelic genome editing by reducing excessive activity of conventional sgRNA-Cas9. Reducing amounts of the all-in-one plasmid or sgRNA-expressing plasmid failed to increase the clones with mono-allelic indels (Fig. 2a and Extended Data Fig. 2). Next, addition of 15-base stretches of guanine [15G], cytosine [15C], adenine [15A], and thymidine [15T] to the 5’ end of spacer sites was tested on the basis of previous reports describing that a few additional guanines at the 5’ end may potentially interrupt the sgRNA-Cas9 activity^23,24^ (Fig. 2b). Importantly, among them, [15C] extension substantially increased frequency of mono-allelic indel clones (Fig. 2c). The [15T]sgRNA almost completely failed to induce indel, which might be due to loss of sgRNA expression because [15T] contained a 4xT transcription termination signal for the U6 promoter^25^. Thus, we focused on cytosine ([C]) extension in the subsequent experiments.

**Fig. 2.**
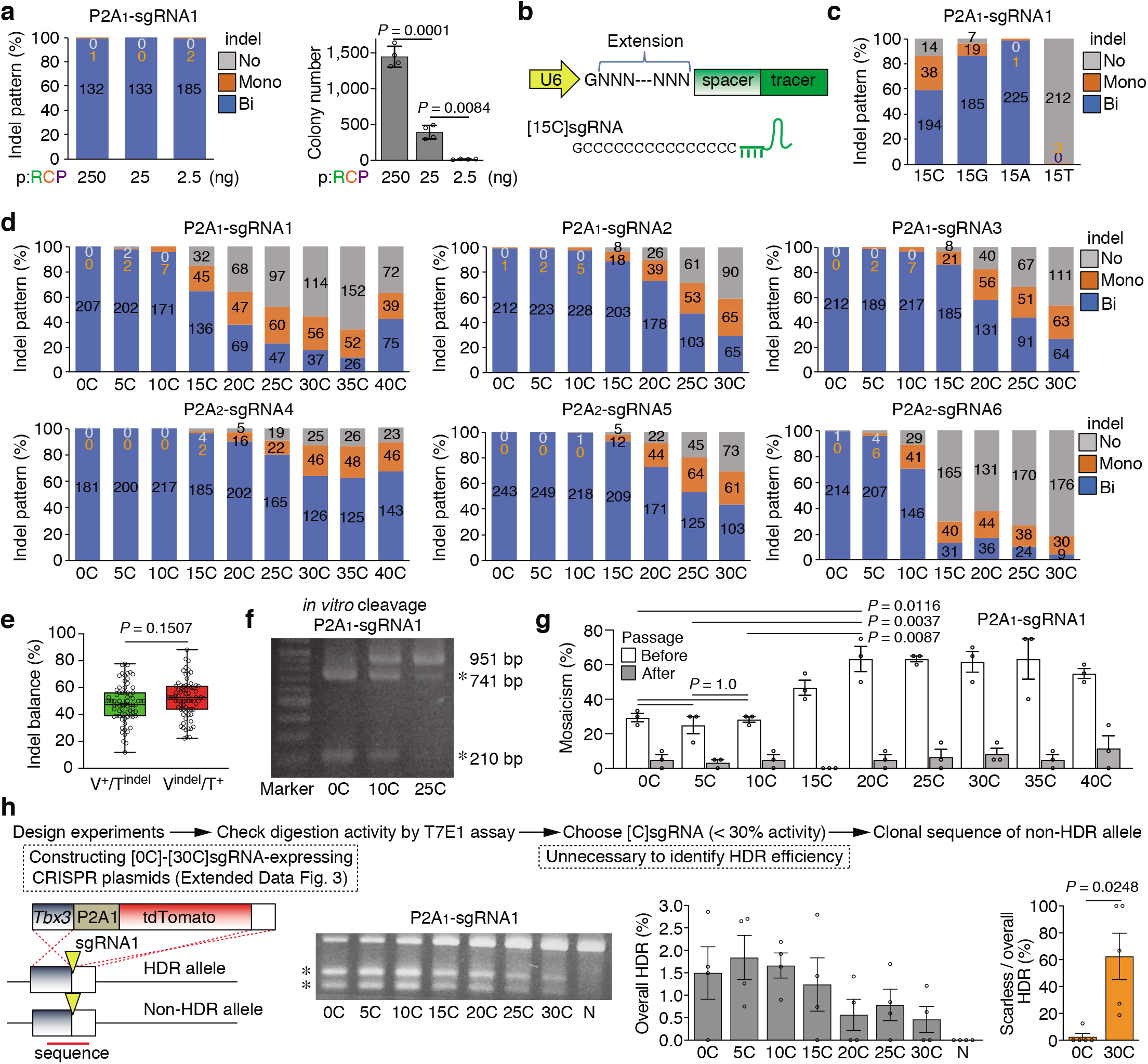
Downsizing of sgRNA-Cas9 activity by addition of 5’-end cytosine stretches. ***a***, Analysis of indel patterns (left) and colony numbers (right) after transfecting all-in-one plasmids containing sgRNA1 at different concentrations in Tbx3-P2A1-AIMS. pRCP, all-in-one plasmid shown in Fig.1d. Data are shown as mean and s.d. from *n* = 4 biological replicates. Total number of clones is shown in each column (also in **c** and **d**). Statistical significance is assessed using Welch’s test with post hoc Games–Howell test. **b**, Schematic of nucleotide extension at the 5’ end of spacer for downsizing sgRNA-Cas9 activity. **c**, Effects of 15-base cytosine [15C], guanine [15G], adenine [15A], and thymidine [15T] extension for sgRNA1 in Cdh1-P2A1-AIMS. Data are shown as mean from *n* = 3 independent experiments performed at different times. **d**, Indel pattern analysis using 6 different sgRNAs in Cdh1-P2A1-AIMS and Cdh1-P2A2-AIMS. Data are shown as mean from *n* = 3 independent experiments performed at different times. **e**, Percentage of mono-allelic indel frequencies for the tdTomato and Venus alleles. Dots indicate individual data points (*n* = 73, 6 sgRNAs, Tbx3- and Cdh1-AIMS). In the box plots with interquartile range, the center lines show medians, and whiskers range from a minimum to a maximum value. Statistical significance is assessed using two-tailed Student’s t-test. **f**, An in vitro cleavage assay to compare [0C], [10C] and [25C]sgRNAs for DSB induction potential. PCR amplicons (951 bp) containing a Tbx3-P2A1-sgRNA1 targeting site is digested by ribonucleoprotein (RNP) composed of [C]sgRNA and SpCas9 recombinant protein. Asterisks indicate digested products (741 bp and 210 bp). **g**, Mosaic frequency before and after passage using different [C]sgRNAs in Cdh1-P2A1-AIMS. Data are shown as mean ± s.e.m. from *n* = 3 independent experiments performed at different times. Statistical significance is assessed using two-way ANOVA and post hoc Tukey–Kramer test. **h**, Induction of mono-allelic HDR without induction of indels on a non-HDR allele. A schematic of HDR of a P2A1-tdTomato cassette at the end of *Tbx3* coding site (left). T7E1 assay to survey indel probability induced by [C]sgRNAs (second from the left). Frequencies of overall HDR (second from the right) and scarless HDR (right) are shown as mean ± s.e.m. from *n* = 4 and *n* = 5 independent experiments, respectively, which are performed at different times. Statistical significance for scarless HDR frequency (right) is assessed using Welch’s t-test. Arrowheads indicate DSB sites. Asterisks indicate PCR products digested by T7E1. N, PX459 plasmid without spacer.

We further investigated the relationships between [C] extension length and bi-/mono-allelic indel patterns by systematically generating all-in-one-plasmids expressing [0C]-[30C]-extended sgRNAs for six different sgRNA sequences (Fig. 2d and Extended Data. Fig. 3, 4). For all six sgRNAs, [C]-extended sgRNAs ([C]sgRNAs) exhibited decreased bi-allelic indels and increased mono-allelic indels in a length-dependent manner, indicating length-dependent editing suppression (Fig. 2d). Allelic bias was not observed in the case of mono-allelic indel induction (Fig. 2e and Extended Data Fig. 5a). Editing efficiency is reportedly influenced by the local genome environment and cell types, even when targeting the same sequences, as confirmed in Extended Data Fig. 5b, c, and highly depends on the target sequences. In fact, the absolute indel probabilities of [C]sgRNAs varied among several different sgRNAs (Fig. 2d and Extended Data Fig. 4a); nevertheless, in all experiments, [C] extension exerted uniform suppression effects on diverse sgRNA sequences. This effect can be explained by the assumption that [C] extension decreases the effective sgRNA-Cas9 complex in a length-dependent manner, as supported by computational modeling (Extended Data Fig. 4b, c and Methods). Accordingly, substantial effect of [C] extension on DSB induction was also confirmed by an *in vitro* cleavage assay using a fixed amount of [C]sgRNA (200 nM), Cas9 (66.7 nM), and template DNA (300 ng) (Fig. 2f). In addition, downsized sgRNA-Cas9 activity by [20C]sgRNAs increased the frequency of mosaicism up to 63 %, while the frequency of mosaicism of conventional system was approximately 30% (Fig. 2g, opened bars). These findings suggest that kinetics of indel induction is delayed by [C] extension and that clonal analysis with dissociation methods is critical for downstream applications, especially when downsized sgRNA-Cas9 is used.

We further investigated whether [C] extension allows both mono-allelic insertion (knock-in, KI) of large gene cassettes via HDR and protection of non-HDR allele from indel induction, i.e. one-step generation of HDR/WT clones (Fig. 2h and Extended Data Fig. 5d-f). Overall HDR frequency, which included HDR/indel clones, gradually decreased along with [C] extension due to the reduction in indel probability (Fig. 2h, middle panels). Although the overall HDR frequency of [30C]sgRNA was 3-fold less than that of [0C]sgRNA, the scarless HDR/WT frequency of [30C]sgRNA was 25-fold higher than that of [0C]sgRNA (Fig. 2h, right panel). These data indicate that one scarless HDR/WT clone can be theoretically obtained by picking 40 or 1.6 tdTomato-positive KI clones when using [0C] or [30C]sgRNA, respectively. Similar result was obtained in AIMS experiment, indicating that one scarless HDR/WT clone can be theoretically obtained by picking 137 or 1.9 G418-resistant KI clones when using [0C] or [25C]sgRNA, respectively (Extended Data Fig. 5d-f). Together, these findings suggest that mono-allelic HDR clones without scar on another allele can be efficiently obtained by downsizing sgRNA-Cas9 activity.

### Computational modeling of single-cell heterogeneity of editing frequency

In theory, maximum of the frequency of mono-allelic indel is 0.5 when setting the indel probably to 0.5 if homogenous cell population was assumed. However, actual frequency of mono-allelic indel (F(Mono)) was substantially lower than estimated F(Mono) especially around the intermediate AIMS[P] levels (AIMS[P] ~ 0.5) (Fig. 3a, b). Thus, we considered heterogeneity in genome editing frequency at the single-cell level and computationally modeled the relationships between [C] extension and mono-allelic indel based on AIMS datasets. By utilizing beta distribution and identifying the optimal setting (α value of 0.715) (Extended Data Fig. 6a-d and Methods), we successfully predicted the frequencies for bi-, mono-, and no-indel that highly matched to the AIMS data (Fig. 3b, c and Extended Data Table 1). The simulation indicated that the highest frequency of mono-allelic indel induction is 30.8% when AIMS[P] is 0.392 (Fig. 3d and Extended Data Table 2). In our experiments, [C] extension between 15 and 30 nucleotides is generally optimal for mono-allelic indel induction (Fig. 2d and Extended Data. Fig. 4). Using this model, we confirmed that both Bac[P] and AIMS[P] yields comparable predictions (Fig.3e) and further observed that Bac[P]-based predictions could be applied even when targeting the endogenous *Alb* gene (Fig. 3f). These results collectively suggest that heterogeneity in editing efficiency is another obstacle for efficient mono-allelic editing and that continuous fine-tuning of Cas9 activity is important to find the optimal range of Cas9 activity.

**Fig. 3.**
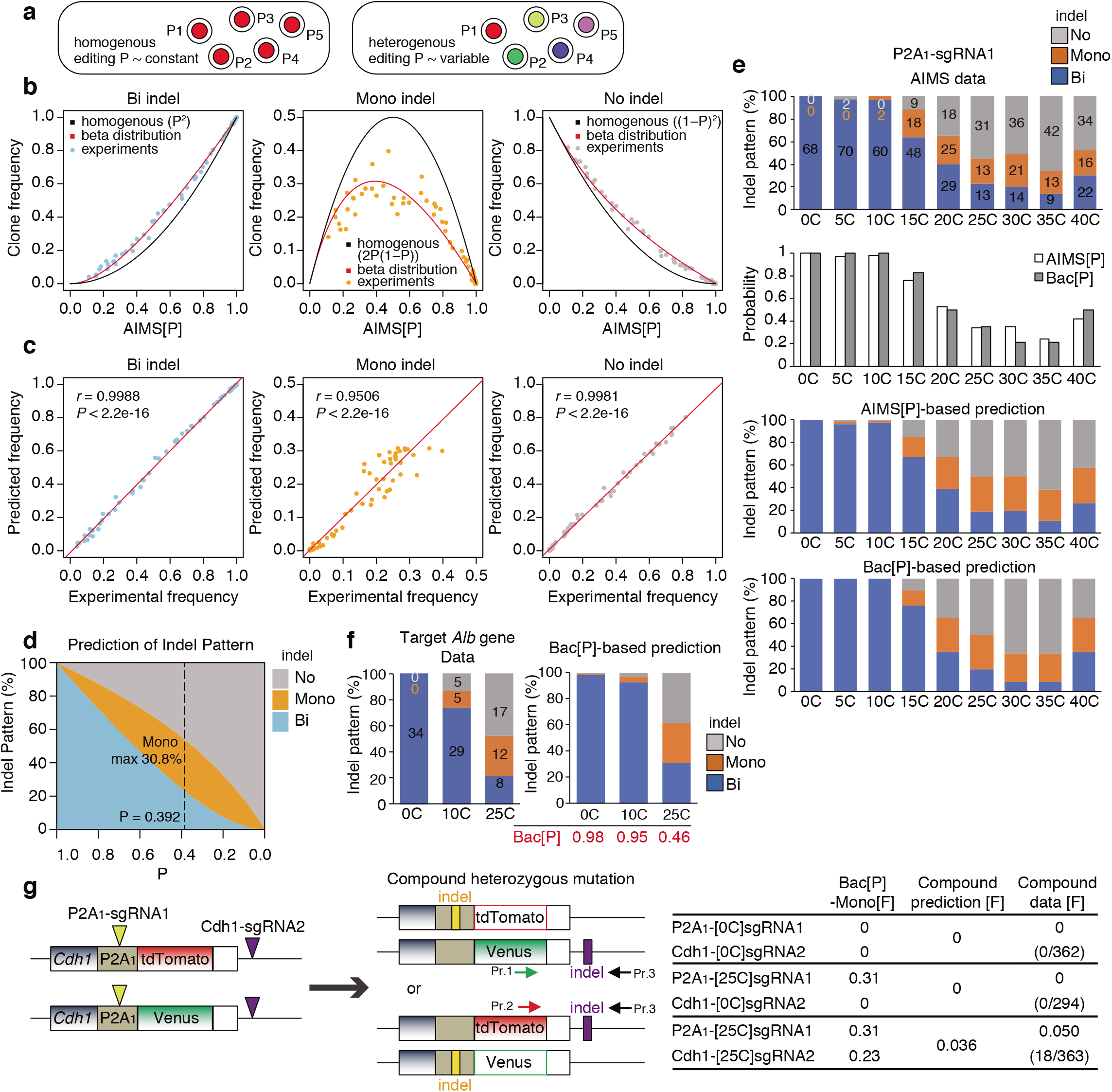
Computational prediction of single-cell heterogeneity of genome editing and bi-/mono-allelic indel frequency. **a**, Homogeneity vs heterogeneity of genome editing frequency at the single cell level. **b**, Relationships between AIMS[P] and clone frequency for Bi-, Mono-, or No indel (*n* = 64). The two curves are generated based on the assumption that single-cell editing probability is homogenous or heterogeneous. In a latter case, beta distribution is adopted. **c**, Correlation between experimental data and beta distribution-based prediction of the frequencies or Bi-, Mono-, or No indel. Linear regression and Pearson’s correlation coefficients (*r*) with P-values are shown. **d**, Relationships between indel probability (P) and allelic indel pattern, simulated by the beta distribution model. The maximum frequency (30.8 %) of mono-allelic indel is obtained when setting P to 0.392. **e**, Comparison between AIMS experimental results with Cdh1-P2A1-sgRNA1 and predictions based on AIMS[P] and Bac[P]. Data are shown as mean from *n* = 3 independent experiments performed at different times and total number of clones analyzed are shown in each column in the AIMS data. **f**, Comparison between experimental data and Bac[P]-based prediction for indel pattern when targeting an endogenous *Alb* (*Albumin*) gene. Data are shown as mean ± s.e.m. from *n* = 3 independent experiments performed at different times and total number of clones analyzed are shown in each column. **g**, Prediction and generation of compound heterozygous mutation clones using Cdh1-P2A1-AIMS. Bac[P]-Mono[F], frequency of mono-allelic indel (Mono[F]) predicted from Bac[P]. Brackets indicate the number of compound heterozygous mutation clones generated by *n* = 3 independent experiments performed at different times. Yellow and purple arrowheads and boxes indicate DSB sites and indels, respectively. Pr.1-3, primers for genotyping.

Next, we examined whether the frequency of compound heterozygous mutation could be predicted using a Cdh1-P2A_1_-AIMS (Fig. 3g). Compound heterozygous clones were obtained only with the [25C]sgRNA combination, and the frequency of 0.50 (18 / 363) was almost identical to the predicted frequency of 0.36 (Fig. 3g), supporting high accuracy of the prediction.

### Generation of a heterozygous SNP disease model

Scarless mono-allelic single nucleotide editing is the most challenging recombination because it involves a high probability of off-target cleavage against a 1 bp mismatched (1mm) HDR allele^9^. We adopted [C] extension method to this issue. To achieve this goal, we chose Fibrodysplasia Ossificans Progressiva (FOP) for which a mono-allelic 617G > A (R206H) mutation in human *ACVR1* gene is a causal mutation^26^ and attempted to generate the identical mutation of mouse *Acvr1* gene in wild-type (WT/WT) mESCs (Fig. 4a). A sgRNA was designed for the region crossing the G>A editing site (Fig. 4a), and indel probability reduction by [C] extension was confirmed by T7E1 and Bac[P] analysis (Fig. 4b, c). After transfection with all-in-one CRISPR plasmids and ssODN as an HDR repair template, sequence analysis was performed for individual clones and the frequencies of overall HDR and precise mono-allelic HDR (WT/R206H) were determined. The overall HDR consists of WT/R206H genotype and all other genotypes harboring indels. [0C]sgRNA induced overall HDR only in 4.1% of clones, however, the frequency of overall HDR for [5C]sgRNA went up to 20.5% and the frequency gradually decreased in parallel with the reduction in indel probability (Fig. 4d). In contrast, the frequency of precise WT/R206H HDR gradually increased with [C] extension, and all clones for [25C] and [30C]sgRNAs exhibited the correct WT/R206H genotype, while [0C]-[10C]sgRNAs could not induce the precise editing (Fig. 4d). Based on overall HDR frequency, we computationally estimated HDR rates after DNA cleavage of single allele by taking into account heterogeneity of single-cell editing efficiency (Fig. 4e). This analysis clearly showed that low HDR rate (2.07 %) increases upon [C] extension and that each [C]sgRNA exhibits similarly high HDR rate (mean 10.99%) except for [25C]. This suggests that [C] extension has generally recovered HDR rate presumably suppressed by conventional CRISPR-Cas9 system.

**Fig. 4.**
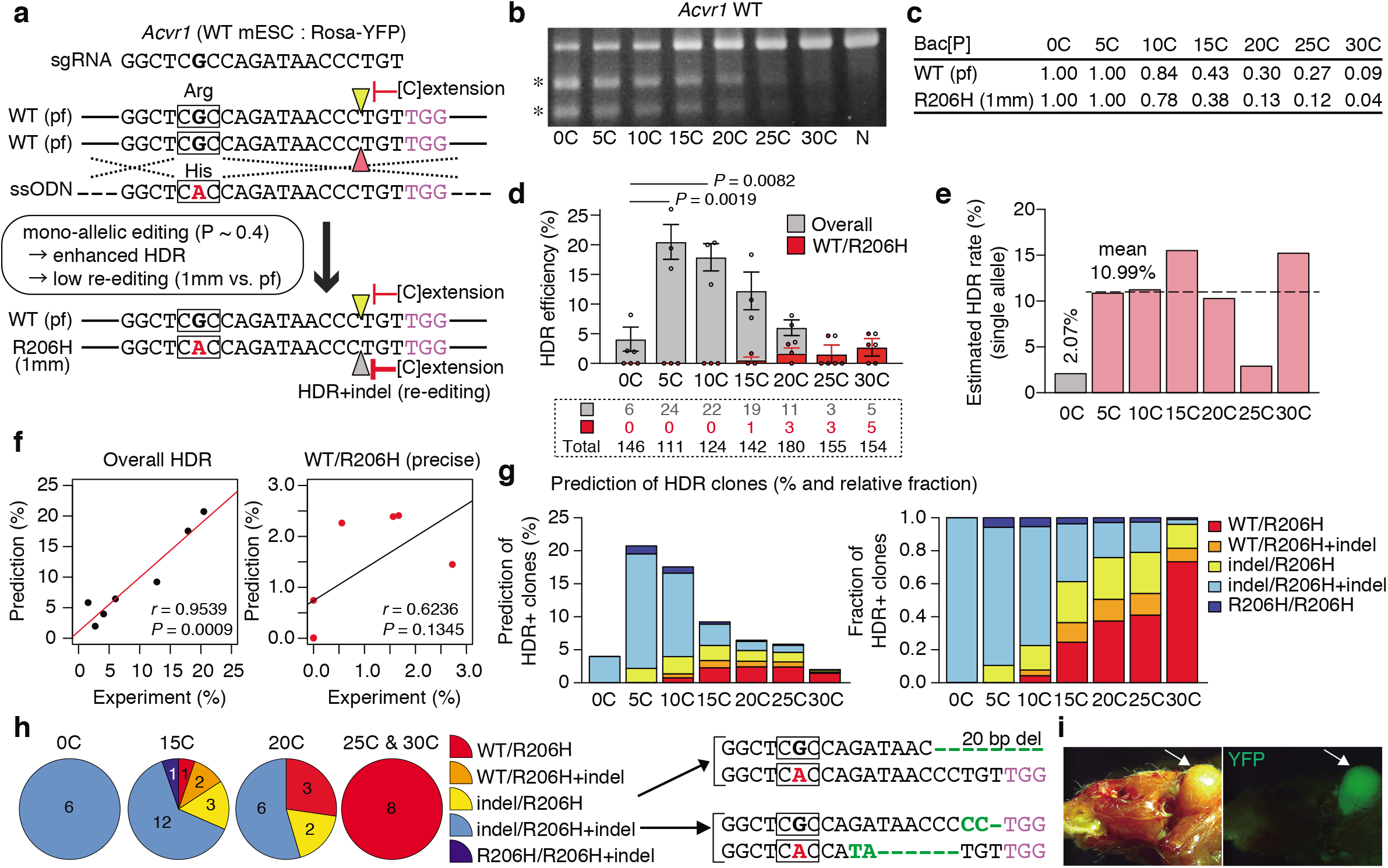
Generation of a heterozygous FOP disease model. ***a***, Schematic of precise HDR for mono-allelic G>A replacement without indels on both HDR and non-HDR alleles in *Acvr1* gene in mESCs. Arrowheads indicate DSB sites. Squares indicate a codon. pf, perfect match; 1mm, 1 bp mismatch. **b**, T7E1 assay. Asterisks indicate PCR products digested by T7E1. N, PX459 plasmid without spacer. **c**, Bac[P] values for both WT and R206H alleles. **d**, Clonal analysis of overall HDR and precise HDR (WT/R206H) efficiencies in the WT/WT mESCs. Overall HDR comprises precise HDR and other HDRs with indels in the HDR- and/or non-HDR-alleles. The number of clones analyzed is shown in a dotted square. Data are shown as mean ± s.e.m. from *n* = 3 independent experiments performed at different times. Statistical significance for overall HDR is assessed using one-way ANOVA and post hoc Tukey–Kramer test. **e**, Computational estimation of HDR rates at single allele. **f**, Correlation between experimental and computationally predicted HDR frequencies. Overall (left) and precise WT/R206H HDR (right) are shown. Linear regression and Pearson’s correlation coefficients (*r*) with P-values are shown. **g**, Prediction of diverse HDR events (left) and relative fraction (right). **h**, Detailed distribution of HDR events shown in the Fig. 4d is shown with actual clone number. Representative sequences of indel/R206H and indel/R206H+indel clones are shown. Squares indicate codon. i, FOP mouse model is generated by microinjection of a WT/R206H clone. An arrow indicates an area of ectopic ossification with mESC contribution, which is traced by the Rosa-YFP reporter.

### Suppression of off-target activity by CRISPR-Cas9 downsizing

In spite of general increase in HDR rates, the precise WT/R206H clones were obtained only for long [C] extension ([20C]-[30C]) but not for short [C] extension with high overall HDR frequency. We assumed that suppressing Cas9 activity makes 1-nucleotide mismatch (1mm) targets less responsive to off-target cleavage, thereby protecting HDR allele from second indel induction. As shown in Fig. 4c, the ratio of off-target editing on R206H HDR allele (1mm) to on-target editing on perfect matched (pf) WT allele decreased along with [C] extension. Since differences in editing efficiency for on-targets and off-targets reflects differences in their dissociation constants, in theory, the ratio of off-target editing to on-target editing and on-target specificity decrease and increase along with suppression of editing frequency, respectively (Extended Data Figure 7a, b and Methods). Thus, protection of HDR allele from second editing becomes remarkable upon long [C] extension. Consistently, the strong off-target inhibitory effect by [C] extension was also confirmed for other sgRNAs in HEK293T cells (Extended Data Fig. 7c, d). We further performed detailed computational modeling of various HDR outcomes based on estimated HDR rates shown in Fig. 4e (Fig. 4f-g, Extended Data Fig. 8a-f, and Methods). The predicted frequencies of overall HDR, WT/R206H HDR, and various HDR patterns were highly consistent with the experimental results (Fig. 4f-h). The optimal indel probability for the precise WT/R206H HDR was predicted to be 0.313, which was slightly lower than the optimal indel probability of 0.392 for mono-allelic indel induction, suggesting the use of [20C]sgRNA and [25C]sgRNA (Extended Data Fig. 8c).

Finally, we confirmed acquisition of the FOP phenotype in the WT/R206H clone by generating chimeric mice. Ectopic ossification at their contributing sites was confirmed (Fig. 4i), as reported in the previous work^27^.

### Safe and systematic precise gene correction in FOP hiPSCs

We next demonstrated the R206H allele-specific gene correction by downsizing sgRNA-Cas9 activity in a FOP patient-derived human induced pluripotent stem cells (hiPSCs, WT/ R206H)^28^ (Fig. 5a). The sgRNA was designed for the R206H (pf) allele and transfected with ssODN containing a silent mutation as a hallmark, which is necessary to distinguish an HDR-corrected (Corrected) allele from original WT allele. Otherwise, WT/- clones, in which PCR amplicons from R206H allele cannot to be obtained due to large deletions or more complex genomic rearrangement^7^, are misidentified as WT/Corrected ones. Efficient indel induction by [0C]sgRNA and its decrease by [5C]-[20C]sgRNAs were confirmed by a T7E1 assay (Fig. 5b). Bac[P] analysis showed that indel probabilities on the WT (1mm) allele decreased with [5C]sgRNA more potently than those on the R206H (pf) allele (Fig. 5c). As described above, the relative suppression of off-target effects is explained by the inherent effects of suppressing on-target activities (Extended Data Fig. 7). Corrected allele (2mm) is further less sensitive to second editing.

**Fig. 5.**
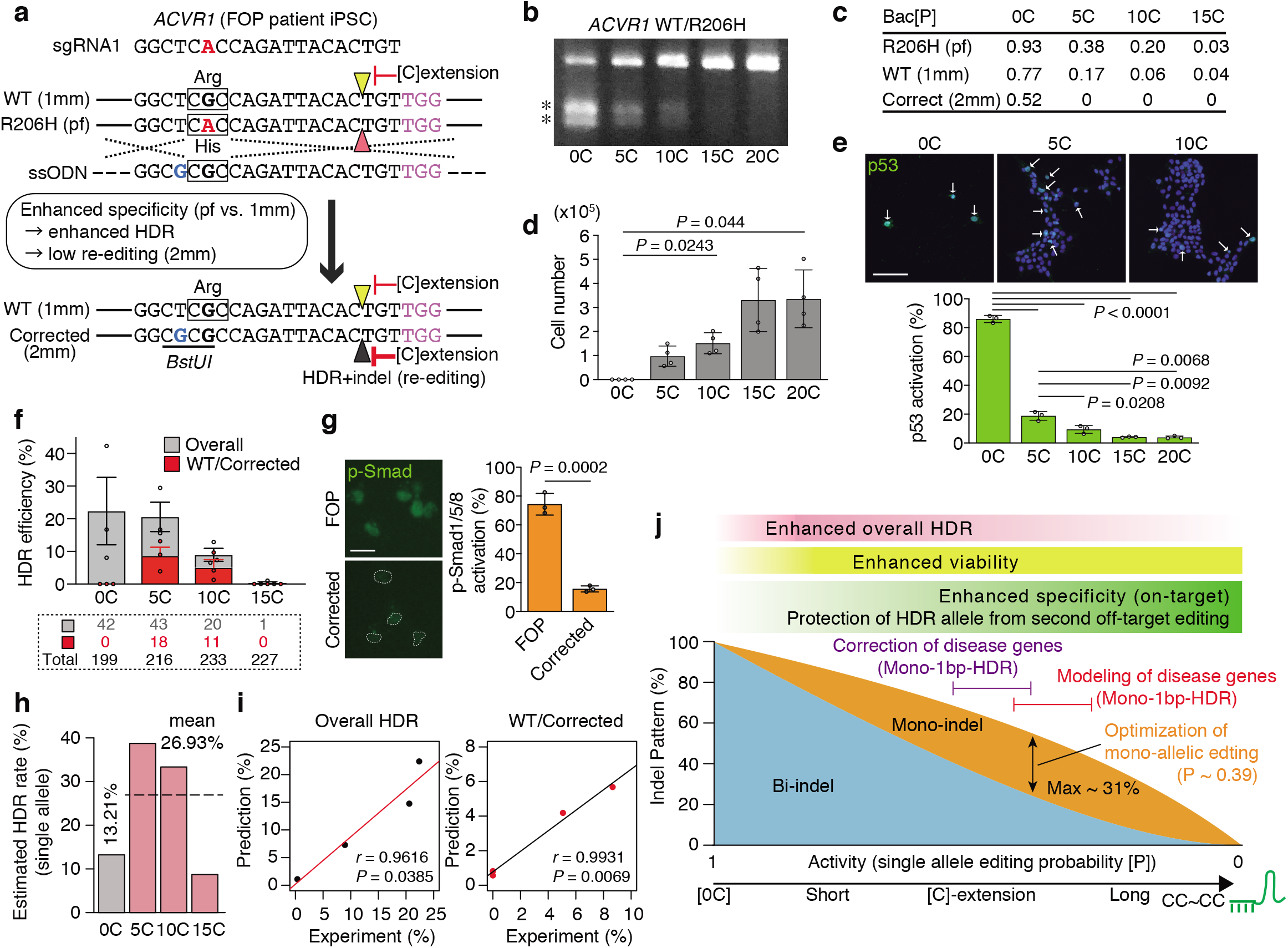
Safe and systematic precise gene correction in FOP hiPSCs. **a**, Schematic of R206H allele-selective precise HDR for A>G correction in FOP iPSCs (WT/R206H). Silent mutation of guanine (G, blue) creates a *BstUI* restriction enzyme site, which allows for rapid selection of HDR clones and for discriminating the Corrected allele (2mm) from original WT one (1mm). Arrowheads indicate DSB sites. Squares indicate codon. pf, perfect match; 1mm, 1 bp mismatch; 2mm, 2 bp mismatches. **b**, T7E1 assay. Asterisks indicate PCR products digested by T7E1. N, PX459 plasmid without spacer. **c**, Bac[P] values for R206H (pf), WT (1mm) and Corrected (2mm) alleles. **d**, Cytotoxicity is examined by counting cell number after all-in-one plasmid transfection and puromycin selection. Data are shown as mean and s.d. from *n* = 4 biological replicates. Statistical significance is assessed using Welch’s test with post hoc Games–Howell test. **e**, Immunocytochemistry to examine p53 activation in the FOP hiPSCs. Data are shown as mean and s.d. from *n* = 3 biological replicates. Statistical significance is assessed using Welch’s test with post hoc Games–Howell test. **f**, Clonal analysis of overall HDR and precise HDR (WT/Corrected) efficiencies in the FOP hiPSCs (WT/R206H). Overall HDR comprises R206- and/or WT-allele-HDR with or without indels. The number of clones analyzed is shown in a dotted square. Data are shown as mean ± s.e.m. from *n* = 3 independent experiments performed at different times. **g**, Immunocytochemistry to examine Activin-induced pSmad1/5/8 activation in the FOP (WT/R206H) hiPSCs and a corrected (WT/Corrected) clone. Data are shown as mean and s.d. from *n* = 3 biological replicates. Statistical significance is assessed using two-tailed Student’s t-test. h, Computational estimation for HDR rate at single allele. **i**, Correlation between experimental HDR frequencies and computational modeling. Overall (left) and precise WT/Corrected HDR (right) are shown. Linear regression and Pearson’s correlation coefficients (*r*) with P-values are shown. **j**, Summary of [C] extension methods.

It is known that Cas9-mediated DSBs induce potent p53-dependent cytotoxicity in hiPSCs^5,6^. Indeed, severe cytotoxicity was induced by [0C]sgRNA (Fig. 5d) and p53 was highly activated in 86% of the surviving cells (Fig. 5e). In contrast, such cytotoxicity and p53 activation were dramatically relieved by the use of [5C]-[20C]sgRNAs (Fig. 5d, e). Inhibition of cytotoxicity by [C] extension was also confirmed by independent experiments targeting other genes in hiPSCs (Extended Data Fig. 9a-d), although HEK293T cells were tolerable to conventional system (Extended Data Fig. 9e-h). We next determined the frequencies of overall HDR and precise WT/Corrected HDR. Overall HDR frequency with [5C]sgRNA was comparable to that of [0C]sgRNA albeit with a lower indel probability, and overall HDR frequency decreased with longer [C] extension (Fig. 5f). The precise WT/Corrected clones could be obtained by [5C]sgRNA and [10C]sgRNA, but not by [0C]sgRNA (Fig. 5f). As the result of gene correction, activin A-mediated activation of bone morphogenetic protein (BMP)-responsive Smad1/5/8 was cancelled in the WT/Corrected clone (Fig. 5g), as reported in the previous work^29^.

We performed similar computational modeling of editing outcomes (Extended Data Fig. 8a, 10a-e). The HDR rates of a single allele for [0C]sgRNA and [5C]-[15C]sgRNAs were estimated to be 13.21% and 26.93%, respectively (Fig. 5h). The predicted overall and WT/R206H HDR frequencies highly correlated with the experimental results (Fig. 5i and Extended Data Fig. 10c). The computational model estimated the frequency of all 12 possible HDR patterns, which suggested that two populations of ‘WT_Corrected_indel/R206H_indel’ (fraction 12) and ‘WT_indel/R206H_Corrected_indel’ (fraction 6) were dominant when indel probability was high (Extended Data Fig. 10e, upper panels), suggesting that lowering indel probability is necessary to prevent second editing and allow single-step precise editing. The optimal indel probability for the precise HDR was simulated to be 0.424, suggesting the use of [5C]sgRNA (Extended Data Fig. 10b, e).

Similar to the mESC data, HDR rate of [0C]sgRNA was estimated to be lower than that of [5-15C]sgRNAs (Fig. 5h). Additional HDR experiment for 3 bp replacement in HEK293T cells also showed HDR enhancement with [5C]sgRNA even though indel probability was similar between [0C] and [5C]sgRNAs (Extended Data Fig. 10f-h). Here we conclude that the precise heterozygous HDR clones can be systematically obtained by downsizing sgRNA-Cas9 activity through multiple mechanisms, including enhancing mono-allelic editing, suppressing p53-dependent cytotoxicity, increasing HDR rates, and suppressing second cleavage of HDR-allele through off-target suppression (Fig. 5j).

## Discussion

Various approaches, such as anti-Cas9 protein and small molecule inhibitor, have been demonstrated to downsize Cas9 activity^14,30–36^. However, their significance in precise genome editing and safety has not been well tested. Here we have established that simple modifications of sgRNAs sufficiently enable fine-tuning of sgRNA-Cas9 activity, which avoid the use of other molecules with unknown adverse effects.

Mono-allelic genome editing using the reduced activity of sgRNA-Cas9 is important not only for generating heterozygous mutants and precise gene correction, but also for protecting the genome from unwanted off-target mutations and cytotoxicity (Fig. 5j). Notably, we clarified an inhibitory effect of [C] extension on the off-target activity on 1mm targets and an enhancement of on-target specificity, although these 1 mm off-targets had the same maximum indel probability as on-targets in the conventional system. Conventional methods inevitably involve multistep editing to generate precise heterozygous clones because HDR-allele have to be protected from second off-target editing by introduction of silent mutations and the silent mutations should be re-corrected especially for non-coding regions^9^. On the other hand, [C] extension methods enable single step heterozygous editing by protecting HDR-allele through direct suppression of off-target effects. This would be beneficial for convenient modeling of heterozygous states of disease mutations and risk variants and investigation of their downstream effects such as allele-specific epigenome and gene regulation. In our system, precise homozygous mutations can be obtained by repeated mono-allelic editing. Our finding is consistent with the context of previous studies suggesting that the increased on-target specificity of engineered Cas9s, truncated gRNAs, and gRNAs with a couple of guanine addition to the 5’ end, is at least partially due to decrease in the activity of sgRNA-Cas9 complex^24,37,38^.

Another important hallmark of [C] extension is suppression of excess DNA damage and cytotoxicity, leading to enhancement of HDR. The previous work reported that p53 activation induces cytotoxicity and inhibits HDR frequency by 19-fold in hiPSCs^6^. We conspicuously demonstrated that even short [C] extension such as [5C]sgRNA had strong potential to reduce p53 activation, enhance cell viability, and HDR rates, while maintaining maximum on-target activity. Thus, to avoid any long-term deleterious effects of excessive DNA damage on cell phenotypes, it may be reasonable to consider the use of sgRNAs with short [C] extension, such as [5C]sgRNA, as the next-generation CRISPR-Cas9 platform for diverse applications in mammalian cells. Furthermore, tunable downsizing of Cas9 activity, presented in this study, offers a systematic and safeguard platform for switchable use of bi-allelic and mono-allelic genome editing.

## Methods

### Cell culture

mESCs were cultured in t2iL medium containing DMEM (Nacalai Tesque) with 2 mM Glutamax (Nacalai Tesque), 1 × non-essential amino acids (NEAA) (Nacalai Tesque), 1 mM Sodium Pyruvate (Nacalai Tesque), 100 U/ml penicillin, 100 μg/ml streptomycin (P/S) (Nacalai Tesque), 0.1 mM 2-mercaptoethanol (SIGMA) and 15% fetal bovine serum (FBS) (GIBCO), supplemented with 0.2 μM PD0325901 (SIGMA), 3 μM CHIR99021 (Cayman) and 1,000 U/ml recombinant mouse LIF (Millipore)^39^. Higher concentration of PD0325901 at 1 μM was also used for 2iL medium. mESC colonies were dissociated with trypsin (Nacalai Tesque) and plated on gelatin-coated dishes. Y-27632 (10 μM, SIGMA) was added when cells were passed. hiPSCs were cultured in mTeSR Plus medium (VERITAS). hiPSC colonies were dissociated with Accutase (Nacalai Tesque) and plated on matrigel-coated dishes (matrigel, CORNING, 3/250 dilution with DMEM). Y-27632 and 1% FBS were added when cells were passed. WT hiPSCs (409B2, HPS0076) were provided by the RIKEN BRC^40^. FOP hiPSCs (HPS0376) were provided by the RIKEN BRC through the National BioResource Project of the MEXT/AMED, Japan^28^. The study using hiPSCs was approved by the Kyushu University Institutional Review Board for Human Genome/Gene Research. HEK293T cells and mouse embryonic fibroblasts (MEFs) were cultured in 10% FBS medium containing DMEM, 2 mM l-glutamine (Nacalai Tesque), 100 U/ml penicillin, 100 μg/ml streptomycin (P/S) (Nacalai Tesque) and 10% FBS. Cells were maintained at 37 °C with 5% CO_2_.

### Animals

C57BL/6 mice (Clea Japan, Tokyo, Japan), ICR mice (Clea Japan, Tokyo, Japan) and *R26R^YFP/YFP^* mice^41^ (a gift from Frank Costantini, Columbia University, New York, NY) were used in the present study. The experiments were approved by the Kyushu University Animal Experiment Committee, and the care and use of the animals were performed in accordance with institutional guidelines.

### Oligo nucleotides

All primers, spacer linkers, and ssODNs used in the present study are listed in the Extended Data Table 3.

### Establishing mESCs

Mouse ES cell lines of B6-5-2 and B6-D2-4 line were established from E3.5 blastocysts of C57BL/6 strain using 2iL or t2iLmedium, respectively, and a mESC line of *R26R^YFP/+^* mouse strain was established using t2iL medium. Blastocysts were placed on feeders (mitomycin C-treated MEFs) after removing zona pellucida. Outgrowths of the inner cell mass (passage number 0, p0) were dissociated with trypsin and plated on gelatin-coated plate (p1). After domed colonies were grown, they were dissociated and passed (p2). The mESC lines were generated by repeating this procedure.

### Generation of AIMS

Knock-in (KI) template plasmids for Cdh1-AIMS were generated by attaching 5’ and 3’ arms to plasmids containing P2A_1_:Venus or P2A_1_:tdTomato cassettes. The P2A_1_ is identical to a P2A sequence which is generally used^18^. The 5’ arm was designed so that the coding end is fused to the P2A sequence in-frame to allow the production of both E-cadherin (CDH1) and the fluorescence protein independently. The KI plasmids for Tbx3-AIMS were constructed using the same strategy. The alternative P2A sequence P2A_2_ was constructed by introducing silent mutations to each codon of the original P2A sequence. Conventional CRISPR-Cas9 system was used to efficiently knock-in the dual color plasmids in a pair of alleles. A spacer linker was designed to induce DSB downstream of the stop codon, and was inserted into *BpiI* sites of a pSpCas9(BB)-2A-Puro (PX459) V2.0 plasmid (Addgene, Plasmid #62988). All sgRNAs used in the present study were designed using CRISPR DESIGN (http://crispr.mit.edu/) or CRISPOR (http://crispor.tefor.net).

Both the constructed all-in-one CRISPR plasmids and dual colored KI plasmids were co-transfected into the mESCs using lipofectamine 3000 (Thermo Fisher Scientific). Dissociated mESCs were plated on gelatin-coated 24-well plates with 500 μl of (t)2iL+Y-27632 medium ((t)2iL+Y). Nucleic acid-Lipofectamine 3000 complexes were prepared according to the standard Lipofectamine 3000 protocol. One μl of Lipofectamine 3000 reagent was added to 25 μl Opti-MEM medium, while 250 ng of each plasmid (all-in-one, Cdh1-P2A-tdTomato, and Cdh1-P2A-Venus plasmid) plus 1 μl of P3000 reagent were mixed with 25 μl of Opti-MEM medium in a different tube. These mixtures were combined and incubated for 5 min at room temperature, and then added to the 24-well plate just after cells were seeded. Twenty-four hours after transfection, puromycin (1.5 or 2 μg/ml) was added for 2 days, and then washed out. The transiently treated-puromycin resistant cells were cultured for several days, and dual color-positive colonies were picked and passed. The genotypes for candidate dual KI clones were confirmed by PCR. Transfection experiments for mouse and human cells were performed according to this procedure, and a couple of passage steps are added when AIMS assay was performed to avoid mosaicism (Fig. 1d). Fluorescent microscopes (BZ-X800, KEYENCE and IX73, OLYMPUS) were used to analyze the AIMS data.

To extract genomic DNA for the clonal sequence analysis, single mESC and hiPSC colonies were suspended in 5-10 μl of 50 mM NaOH (Nacalai Tesque) and incubated at 99 °C for 10 minutes.

### Plasmid construction

To generate all-in-one CRISPR plasmids for [5C](3A), [10C](8A), [15C](13A), [20C](18C), [25C](23A) and [30C](28A)sgRNA expression, spacer linkers were inserted into the *BpiI* sites of a PX459 plasmid (Extended Data Fig. 3). In the plasmids, 3^rd^, 8^th^, 13^th^, 18^th^, 23^rd^ or 28^th^ cytosine was replaced with adenine since overhang sequence of CACC is required for linker ligation. The standard spacer linkers were inserted into the *BpiI* sites of the [5C](3A), [10C](8A), [15C](13A), [20C](18A), [25C](23A) or [30C](28A) PX459 plasmid, leading to generation of [5C]-[30C]sgRNA expressing all-in-one plasmids.

For a plasmid dilution assay, sgRNA expressing plasmid was constructed by removing a Cas9-T2A-Puro cassette from a PX459 plasmid using *KpnI* and *NotI* sites. The different amount of sgRNA expressing plasmid (0-250 ng) was co-transfected with unmodified PX459 plasmid (250 ng).

### *In vitro* DNA cleavage assay

For preparation of template DNA to be cleaved, 951 bp fragment was amplified by PCR using a Tbx3-P2A_1_-Venus KI plasmid. The [0C], [10C] and [25C]sgRNAs were synthesized by *in vitro* transcription (IVT) using a T7 RiboMAX Express Large Scale RNA Production System (Promega) according to the manufacture’s protocol. The template DNA fragments required for IVT were amplified by PCR using forward and reverse primers. The T7 promoter sequence and cytosine tails were added to 5’ end of the forward primer. Cas9 protein (IDT) was suspended with a Diluent B (NEB) to make 1 μM solution. The 10 × Cas9 reaction buffer contains 1 M HEPES, 3 M NaCl, 1 M MgCl_2_, and 250 mM EDTA (pH6.5). For cleavage, 1 μl of Cas9 (1μM), 1μl of [C]sgRNAs (3 μM), 1 μl of 10 × Cas9 reaction buffer, and 3 μl of template DNA (100 ng/μl) were mixed with 9 μl of distilled water (Total 15 μl reaction volume) and reacted at 37 °C for 1 hour.

### Scarless mono-allelic KI of tdTomato or P2A_1_-Neo cassette

A Tbx3-P2A_1_-tdTomato KI plasmid was co-transfected with Tbx3-sgRNA1-expressing PX459 to the mESCs. After transient puromycin selection, colonies were passed after dissociation and the subsequent colonies were analyzed. The colonies with mosaic tdTomato expression were excluded for the data analysis. After counting the colony number, tdTomato positive colonies were picked and genomic DNA was extracted for sequencing.

Neomycin (Neo) KI plasmid was constructed by replacing tdTomato cassette of the Tbx3-P2A_1_-tdTomato KI plasmid with a P2A_1_-Neo cassette. The KI plasmid was co-transfected with P2A_1_ sgRNA1-expressing PX459 to a Tbx3-P2A_1_-AIMS clone. When puromycin was removed, geneticin (400 μg/ml, GIBCO) were added to select KI clones. As analyzed all 8 clones were confirmed to have successful KI genotypes, geneticin-resistant colonies were counted as KI.

### T7E1 assays

PCR reactions to amplify specific on-target or off-target sites were performed using KOD-Plus-ver.2 DNA polymerase (TOYOBO) according to the manufacture’s protocol. Resulting PCR amplicons were denatured and re-annealed in 1 × NEB buffer 2 (NEB) in a total volume of 9 μl using a following conditions: 95 °C for 5 min; 95 °C to 25 °C ramping at −0.1 °C /s, and hold at 4 °C. After re-annealing, 1 μl of T7 Endonuclease I (NEB, 10 units/μl) was added and incubated at 37 °C for 15 min.

### Bac[P] assays

Purified PCR products which amplified specific on-target or off-target sites were inserted into a T-easy vector (Promega) and transformed into DH5-α bacterial cells. To enable rapid and efficient indel detection, plasmids were directly isolated from each white colony (blue white screening was done), and inserted DNA fragment was amplified by PCR. The PCR amplicons were mixed with the PCR products which were amplified from wild-type DNA template such as KI plasmids or unedited genomic DNA, and T7E1 assay was performed. Sanger sequencing was also performed for the PCR amplicons, which were not digested by T7E1, to determine the total number of colony harboring an indel. The Bac[P] value was calculated by the following formula: Bac[P] = *Indel* /*Total*.

Bac[P] values for both WT and R206H alleles were determined with the experiments inducing indels using various [C]sgRNAs in the mESC clone of the FOP model. The targeting sites of both WT and R206H alleles were amplified by PCR and were cloned into T-easy vector. Sanger sequencing was performed for each PCR product derived from single bacterial clones as described above. Similarly, Bac[P] values for both R206H (pf) and WT (1 mm) alleles were determined by inducing indels in the FOP hiPSCs, while a corrected cell line (WT/Corrected) was used to determine Bac[P] value of the corrected allele (2 mm). Since some PCR products do not contain a G/A hallmark due to intermediate-sized deletions from 12 to about 50 nucleotides, it is not possible to determine which allele was edited for these PCR products. We observed that the fraction of such products with intermediate-sized deletions was relatively constant (approximately 20 % in the experiments in Fig. 4 and 10~20 % in the experiments in Fig. 5) and did not decrease along [C] extension, suggesting that the intermediate-sized deletions are byproducts of short indel induction processes. We therefore assigned to the products with intermediate-sized deletions to two alleles using the ratio of PCR products whose origins were convincingly confirmed. For the analysis in Fig. 5, averages of Bac[P] for WT (1 mm) allele based on comparisons of (1) R206H (pf) and WT (1 mm) alleles and (2) WT (1 mm) and corrected (2 mm) alleles were used for the subsequent computational analysis.

### Cellular viability assays

Based on the transfection protocol described above (‘Generation of AIMS’), 2 × 10^5^ WT hiPSCs or 4 × 10^4^ HEK293T cells were seeded on the 48-well plate and transfected with 100 ng of the all-in-one CRISPR plasmids (2/5 scale of 24-well plate version). The hiPSCs cells were dissociated and counted using trypan blue 3 or 4 days after the transient puromycin treatment at 1.5 μg/ml, while HEK293T cells were counted 4 days after the transient puromycin treatment at 3 μg/ml.

### Generation and correction of a FOP model via HDR with ssODNs

The transfection protocol for 24-well plate experiment is described above (‘Generation of AIMS’). For HDR induction in mESCs and HEK293T cells, 1 μl of 10 μM ssODN (eurofins) was added to the plasmid-lipofectamine complex, whereas 1 μl of 3 μM ssODN was added when hiPSCs were transfected since 10 μM concentration induced severe toxicity. After transient puromycin selection, colonies were dissociated and plated at low density to avoid mosaicism. Single colonies were picked and genomic DNA was extracted. Sequence analysis was performed to identify G to A replacement with or without indel. For the correction of the FOP hiPSCs, clones receiving HDR were screened by digesting PCR product using *BstUI* restriction enzyme (NEB), and then the *BstUI*-positive PCR products were sequenced.

### Immunocytochemical analysis

For p53 staining, transfection for HDR induction was performed in 1/5 scale of a 24-well plate experiment, according to the protocol described above. In this assay, 6 × 10^4^ hiPSCs were seeded on a matrigel-coated 96-well plate in triplicate. Puromycin selection was performed to examine p53 activity only in the transfected cells. The survived cells were fixed with 4% PFA two days after puromycin removal. For pSmad1/5/8 staining, 5 × 10^3^ cells were plated on a matrigel-coated 96-well plate without Y-27632 and 1% FBS. After 2.5 h culture, Activin-A (100 ng/ml) (R&D Systems) was treated for 30 min, and the cells were fixed with 4% PFA. Antibody reaction was performed following standard protocols. Rabbit polyclonal p53 (FL-393, Santa Cruz, 1:200) and rabbit monoclonal pSmad1/5/8 (D5B10, Cell Signaling Technology, 1:1000) antibodies were reacted overnight at 4 °C. A donkey anti-rabbit Alexa Fluor 488 secondary antibody (Thermo Scientific, 1:1000) was reacted RT for 30 min. Data analysis was performed by a cell count application of a fluorescent microscope to select cells with p53 and pSmad1/5/8 activation by setting fluorescence intensity thresholds (BZ-X800, KEYENCE).

### Chimera generation

A mESC clone of a FOP model (C57BL/6 strain) was dissociated with trypsin and 5-8 cells were injected into the 8-cell embryos (E2.5) harvested from ICR pregnant mice. Injected blastocysts were transferred into the uteri of pseudo-pregnant ICR mice. Chimeric contribution was confirmed by coat-color and YFP fluorescence. YFP was observed using a fluorescence stereo microscope (M165FC, Leica).

### Computational modeling and analysis of single-cell level genome editing, [C] extension, and HDR efficiency

#### Comparison of AIMS[P] and Bac[P]

In our study, the probability of single allele editing (P) is determined by two ways: (1) AIMS and (2) Bac[P] assay based on T7E1 assay and complementation by sequence validation. AIMS-based P (AIMS[P]) was determined as:

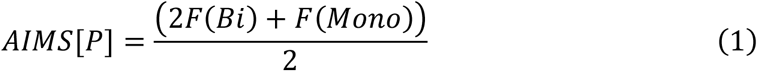
where F(Bi) and F(Mono) are the experimental frequency of cells with bi-allelic and mono-allelic genome editing.

In the initial phase of the study, we compared matched AIMS[P] and Bac[P] for nine sgRNAs (Cdh1-P2A_1_-sgRNA1 with different lengths of [C] extension) and observed that AIMS[P] correlated well with Bac[P] (Extended Data Fig. 6a). In the subsequent analyses, we used AIMS[P] for modeling of indel insertion frequency in Figures 2 and 3 and Bac[P] for modeling of HDR frequency in Figures 4 and 5.

#### Modeling of genome editing frequency at the single-cell level

We performed extensive analysis by combining AIMS and generation of sgRNAs with various [C] extension. When the editing efficiency is homogenous across the cell population, the frequency of cells with bi-allelic, mono-allelic or no genome editing, F(Bi), F(Mono), or F(No) can be estimated as:

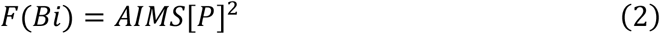

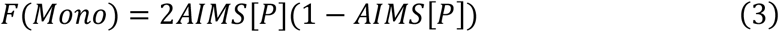

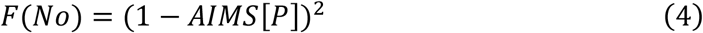

We first used these equations and observed that actual F(Mono) is lower than estimated F(Mono) especially around the intermediate AIMS[P] levels (AIMS[P] ~ 0.5). Thus, heterogeneity in genome editing frequency at the single-cell level was considered and modeled using beta distribution. Probability density functions of P and mean P (E(P)) are given by:

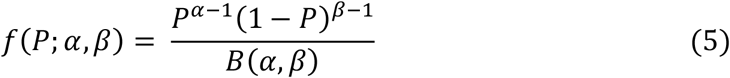

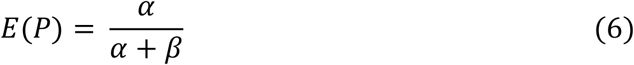

Mean P corresponds to AIMS[P] (or Bac[P]). Using beta distribution, F(Bi), F(Mono), F(No) can be described as:

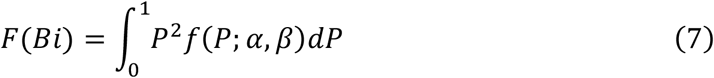

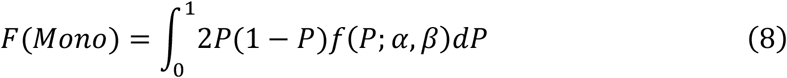

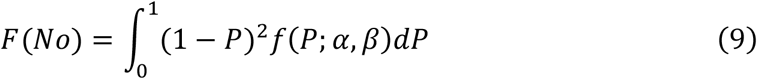

Using these equations, we first determined α value for each experiment that minimized squared residuals between experimental F(Bi), F(Mono), and F(No) and simulated F(Bi), F(Mono), and F(No) (Extended Data Fig. 6b). As shown in Extended Data Fig. 6b, we observed that optimized α values were relatively constant in a wide range of AIMS[P] (0.1 < AIMS[P] < 0.9). Thus, we next used the sum of squared residuals (SSR) as the error function: SSR = Σ(Experimental data – Simulated data)^2^, and determined a constant α value that minimized SSR (Extended Data Fig. 6c, left, *α* = 0.715). Probability density functions with different mean P are shown in a middle panel of Extended Data Fig. 6c. We confirmed that introduction of beta distribution greatly reduced SSR compared to the setting of homogenous editing frequency (Extended Data Fig. 6c, right) and well explained the experimental F(Bi), F(Mono), and F(No) along diverse AIMS[P] (Extended Data Fig. 6d). In addition, we tested normal distribution to approximate the heterogeneity in genome editing frequency at the single-cell level, but observed that beta distribution was superior to normal distribution.

#### Effects of [C] extension on CRISPR-Cas9 system

The efficiency of the single-allele editing P (P(pf); pf: perfect match) can be described as:

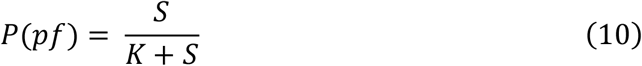
where the concentration of effective sgRNA-Cas9 complexes and the dissociation constant between the sgRNA and its target site are defined as S and K, respectively. Based on high editing efficiency without [C] extension (P ~ 1), we assumed that the recovery rate from single-site damage is very low and did not consider in subsequent analysis. To mechanistically understand the effects of [C] extension and 1 mismatch, we further assumed that [C] extension and 1mismatch decreases S and increases K, respectively. By setting S to 1 for each of different sgRNA sequences without [C] extension, we first approximated K values for each of different sgRNA sequences (8 sgRNA sequences). When P (AIMS[P] or Bac[P]) was 1, P was set to 0.99. Next, relative S concentrations were determined using K and AIMS[P] for sgRNAs with [C] extension. While the relationships between [C] extension and AIMS[P] varied among different sgRNA sequences (Extended Data Fig. 4a), we found clear and similar inverse relationships between [C] extension and relative S values for different sgRNA sequences (Extended Data Fig. 4b). A linear regression gave a good fit to logarithm of relative S against the length of [C] extension for all sgRNA sequences (Extended Data Fig. 4b). Furthermore, analysis of covariance (ANCOVA) indicated that slopes of linear regression did not significantly differ among various sgRNA sequences (Extended Data Fig. 4c). This suggests that [C] extension exerts uniform suppression effects on diverse sgRNA sequences.

#### Effect of cytosine extension on specificity of CRISPR-Cas9 system

As described above, 1mm (or 2mm) is considered to increase K in the equation (10). The efficiency of the single-gene editing P on 1mm (or 2mm) target can be described as:

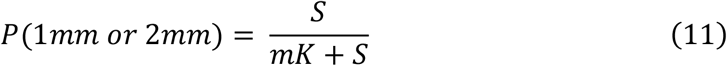
where m is the ratio of K for 1mm target to K for perfect match target.

Thus, the single-gene editing P on 1mm (or 2mm) can be expressed as the function of P(pf) as:

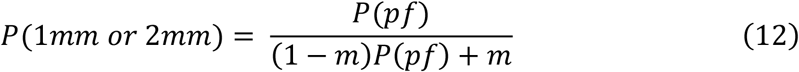

For the results in Figures 4 and 5, we determined m that fits to matched P(pf) and P(1mm or 2mm) by using SSR as the error function (Extended Data Fig. 7a).

The ratios between P(pf) vs. P(1mm or 2mm) are also described as the function of P(pf) as:

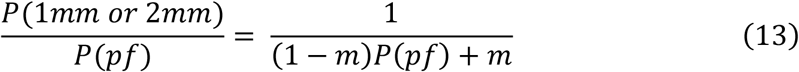

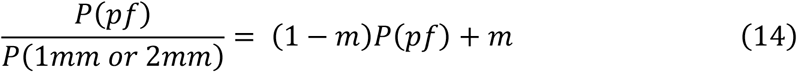

As shown in Extended Data Fig. 7b, decreasing P(pf) contributes to reduction of relative off-target ratio and increases in specificity. Thus, downsizing CRISPR-Cas9 activities by [C] extension is also beneficial for reduction of relative off-target activities and enhancement in specificity.

#### Modeling of HDR frequency from homozygous states (Figure 4)

Using beta distribution, frequencies of the various HDR clones in Figure 4 are determined as follows (Extended Data Fig. 8a, b):

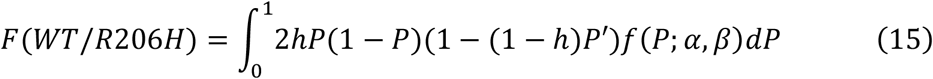

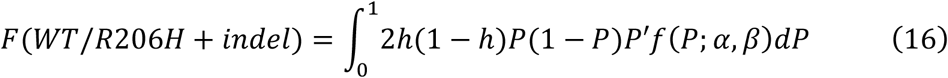

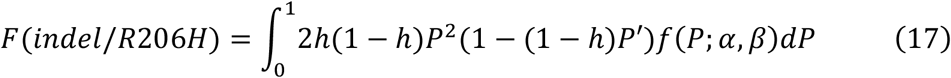

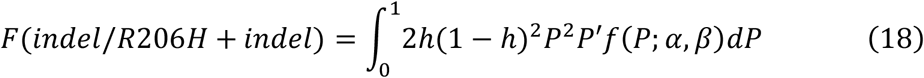

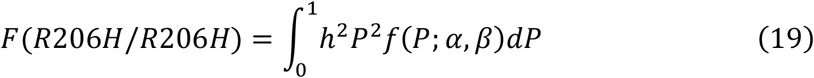

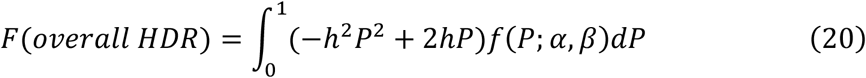
where efficiency of HDR on Cas9-cleaved single allele and the probability of the single-gene editing on edited target, i.e. 1mm target, are defined as h and P’, respectively (Extended Data Fig. 8b). P’ is described in the same manner to equation (12):

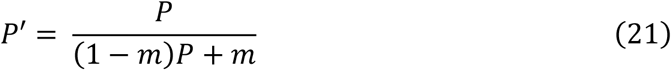
where m is 1.723. [C] extension decreases P according to the length of [C] extension (Extended Data. Fig. 8c).

For simplicity, h was considered to be constant across cell population for each experiment. Based on the experimental results of overall HDR frequencies and the equation (eq. #20), h was estimated for each [C] extended sgRNA (Fig. 4e). While h without [C] extension was very low (2.07%), h with [C] extension was generally high around 11%. This suggests that conventional system without [C] extension reduces HDR and [C] extension releases the suppression to the upper limit of HDR. Based on these findings, we used mean of estimated h (10.99%) as the h for [C] extended sgRNAs and estimated the frequency of distinct HDR patterns, overall HDR and precise HDR (Fig. 4f, g). For sgRNA without [C] extension, estimated h (2.07%) was used. The simulated data well fitted to the experimental results (Fig. 4f, g). To predict HDR outcomes continuously, we designed hypothetical function of h along P (h = 2.07% (P > 0.9); h = 10.99% (P < 0.9)) (Extended Data Fig. 8d) and estimated the frequency of distinct HDR patterns, overall HDR and precise HDR (Extended Data Fig. 8e, f). In the simulation, maximum of precise HDR is obtained when P = 0.313 (Extended Data. Fig. 8c, f).

#### Modeling of HDR-based gene correction (Figure 5)

Using beta distribution, frequencies of the various HDR clones in Figure 5 are determined as follows (Extended Data Fig. 8a and 10a, b):

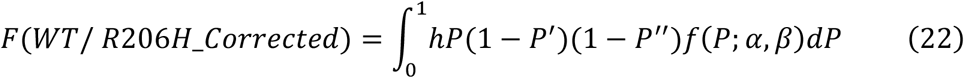

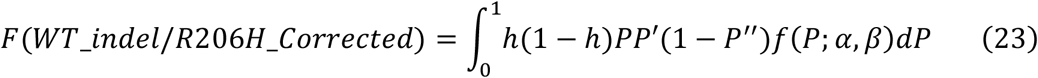

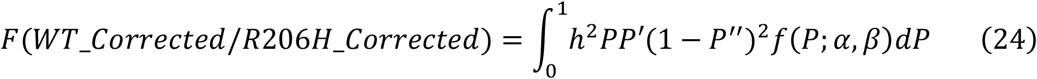

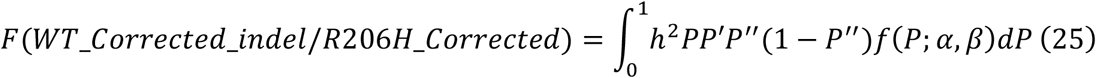

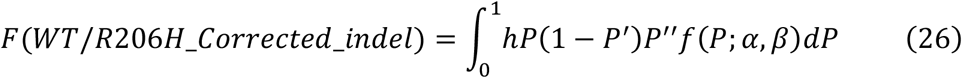

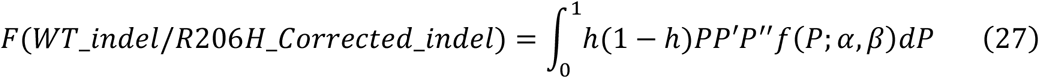

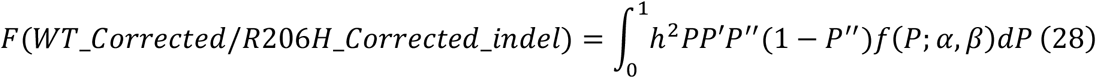

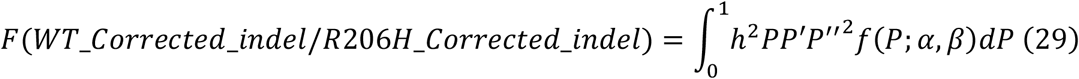

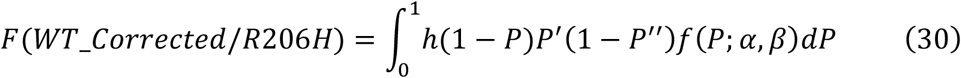

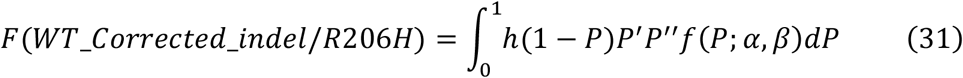

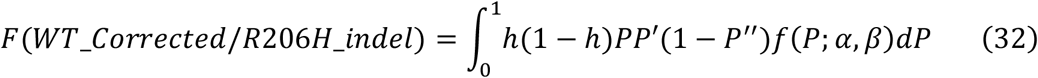

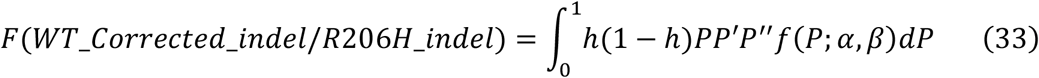

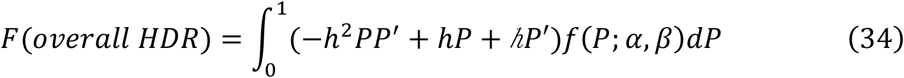
where efficiency of HDR on Cas9-cleaved single allele and the probability of the single-gene editing on WT or HDR-corrected target, i.e. 1mm or 2mm target, are defined as h and P’ or P’’, respectively (Extended Data Fig. 10a). P’ and P’’ is described in the same manner to equation (12):

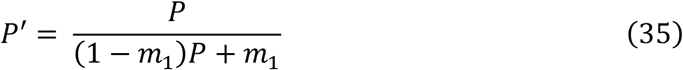

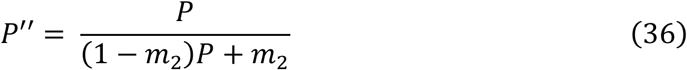
where m1 is 3.459 and m2 is 12.0793, respectively (Extended Data Fig. 7). The relationship between [C] extension and P is shown in Fig. 5c and Extended Data. Fig. 10b.

For simplicity, h was considered to be constant across cell population for each experiment. In addition, HDR rate on R206H and WT allele was considered to be same. Based on the experimental results of overall HDR frequencies and the equation (eq. #34), h was estimated for each [C] extended sgRNA (Fig. 5h). Consistent with the results in Fig. 4, h with [C] extension was higher than h without [C] extension. Together with the results in Fig. 4, this suggests that conventional system without [C] extension reduces HDR probably due to extensive DNA damage and p53 response (as examined in Fig. 5d, e) and [C] extension releases the suppression to the upper limit of HDR. We also observed that h in Fig. 5 was generally higher than h in Fig. 4. This may be because cell lines used in Fig. 5 have only one perfect match target, thus eliciting weaker suppressive effects on HDR rate, while cell lines used in Fig. 4 have two perfect match targets. Based on these findings, we used mean of estimated h (26.93%) as the h for [C] extended sgRNAs and estimated the frequency of overall HDR and precise HDR (Fig. 5i and Extended Data Fig. 10c). For sgRNA without [C] extension, estimated h (13.21%) was used. The simulated data well fitted to the experimental results (Fig. 5i and Extended Data Fig. 10c). To predict HDR outcomes continuously, we designed hypothetical function of h along P (h = 13.21% (P > 0.9); h = 26.93% (P < 0.9)) (Extended Data Fig. 10d) and estimated the frequency of distinct HDR patterns, overall HDR and precise HDR (Extended Data Fig. 10e). In the simulation, maximum of precise HDR is obtained when P = 0.424 (Extended Data. Fig. 10b, e).

#### Statistics

Sample sizes were determined based on our previous experience of performing similar sets of experiments. Statistical tests were performed using JMP (14.2.0) and R (3.2.1). We verified the equality of variance assumption by using the F-test or Levene test. As a pre-test of normality, we used the Kolmogorov–Smirnov test. Differences between two groups were analyzed using two-tailed Student’s t-test (Fig. 2e) or Welch’s t-test (Fig. 2h, right). Comparisons among more than two groups were analyzed using one-way or two-way ANOVA and post hoc Tukey–Kramer test (Fig. 2g, 4d, and Extended Data Fig. 5b, e, f) or Welch’s test with post hoc Games–Howell test (Fig. 1g, 2a, 5d, e, and Extended Data Fig. 9b, d, f, h). In all bar graphs, data are expressed as mean ± s.e.m. (Fig. 2g, h, 4d, 5f, and Extended Data Fig. 1d, 5a-c, e, f) or ± s.d. (Fig. 2a, 5d, e, g, and Extended Data Fig. 9b, d, f and h). In the scatter dot plots, center lines show medians; whiskers show 25th and 75th percentiles from median (Fig. 1g). In the box plots, center lines show medians; box limits indicate the 25th and 75th percentiles; whiskers range from a minimum to a maximum value (Fig. 2e).

#### Data availability

All data will be made available upon request to the corresponding author.

#### Code availability

Scripts needed to reconstruct analysis files will be made available on request.

## ACKNOWLEDGEMENTS

We thank Dr. Frank Costantini for providing *R26R^EYFP/YFP^* mice; Atsushi Miyawaki and Hiroyuki Miyoshi for sharing reagents; Yuuki Honda, Mariko Tasai, Chiaki Kaieda, Shusaku Abe, Takashi Ishiuchi, and Hiroyuki Sasaki for excellent technical assistance. This work was supported in part by grants from the JSPS KAKENHI (17K15046, 18H04737, and 20H05041 to M.K.; 19K24694 to H.I.S.; 18H05102, 19H01177, 19H05267, and 20H05040 to A.S.), the AMED (JP20ae0201012h to M.K.; JP20bm0704034 to A.S.), Center for Clinical and Translational Research of Kyushu University Hospital (to M.K.), Takeda Science Foundation (to M.K. and A.S.), Uehara Memorial Foundation (to A.S.), and Mitsubishi Foundation (to A.S.). Studies conducted by H.I.S. at the Massachusetts Institute of Technology were supported by United States Public Health Service grants R01-GM034277 and R01-CA133404 to P. A. Sharp and P01-CA042063 to T. Jacks from the NIH, as well as a Koch Institute Support (core) grant P30-CA14051 from the National Cancer Institute, and supported in part by an agreement between the Whitehead Institute for Biomedical Research and Novo Nordisk.

## AUTHOR CONTRIBUTIONS

M.K. conceived and designed the research. M.K. and R.K. performed experiments. M.K and H.I.S. analyzed data. H.I.S performed computational analysis. M.K., H.I.S. and A.S. wrote the manuscript. All authors read and approved the final manuscript. A.S. supervised the project.

---

**Extended Data Fig. 1.**
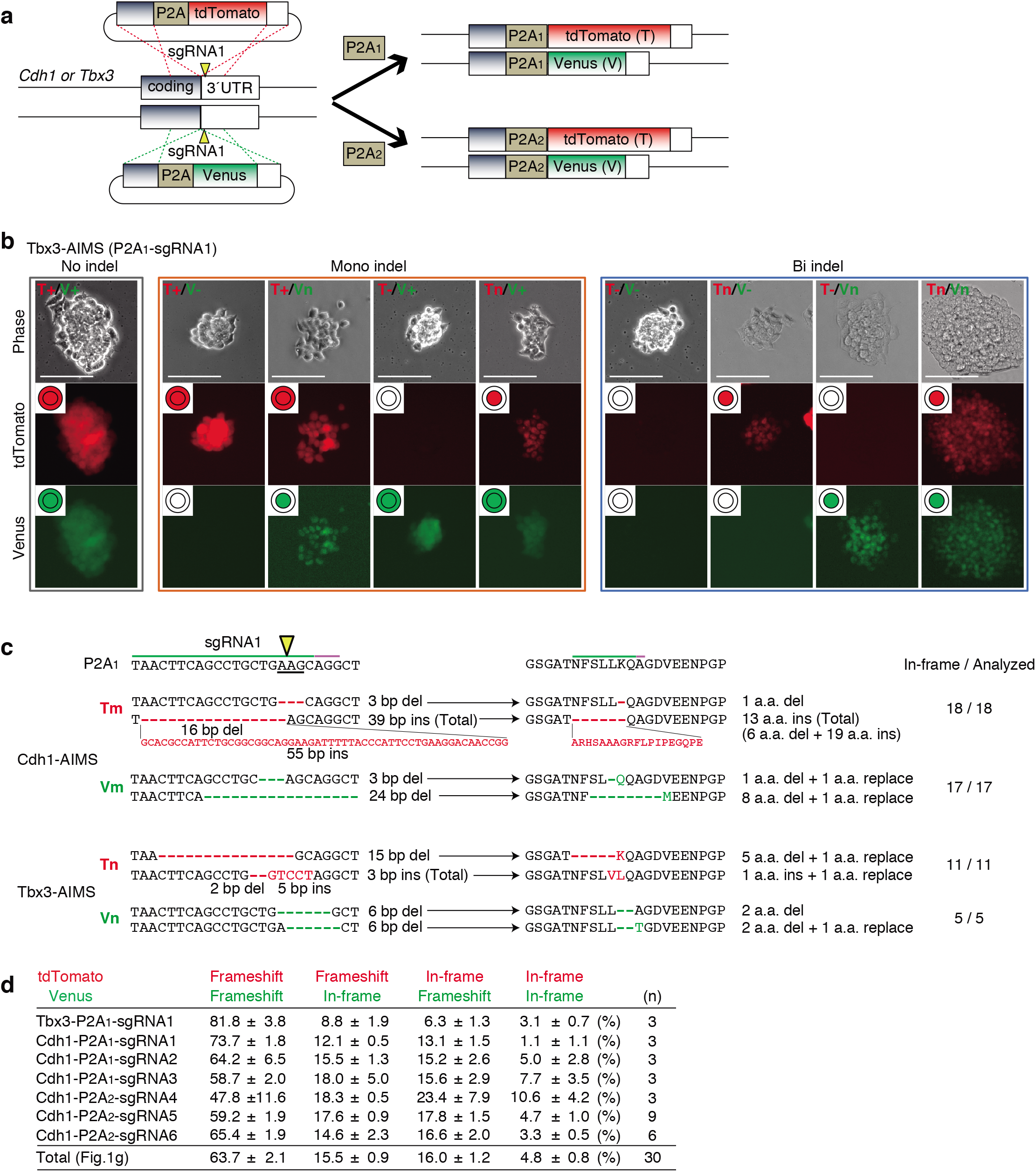
AIMS construction and indel analysis. **a**, Schematic of generating dual color KI mESC clones for AIMS. Two types of targeting plasmids are simultaneously knocked into the two alleles of the *Cdh1* or *Tbx3* locus using CRISPR-Cas9. Arrowheads indicate DSB sites. **b**, Representative results of Tbx3-P2A1-AIMS. Genotypes are determined by nine combinations of tdTomato/Venus expression and localization in Tbx3-P2A1-AIMS. T, tdTomato; V, Venus; +, no indel represented by both cytosolic and nuclear localization; n, in-frame indel represented by nuclear localization; -, frameshift indel or large deletion represented by loss of fluorescence. Scale bar indicates 100 μm. **c**, Representative DNA and amino acid sequences of in-frame indels. The clones showing cytosolic expression (T+ or V+) are sequenced and the number of clones with in-frame indel are shown. **d**, The table shows percentages of the four types of bi-allelic indel patterns. Total indicates the mean of all data (*n* = 30), which is shown in Fig. 1g. Data are shown as mean ± s.e.m..

**Extended Data Fig. 2.**
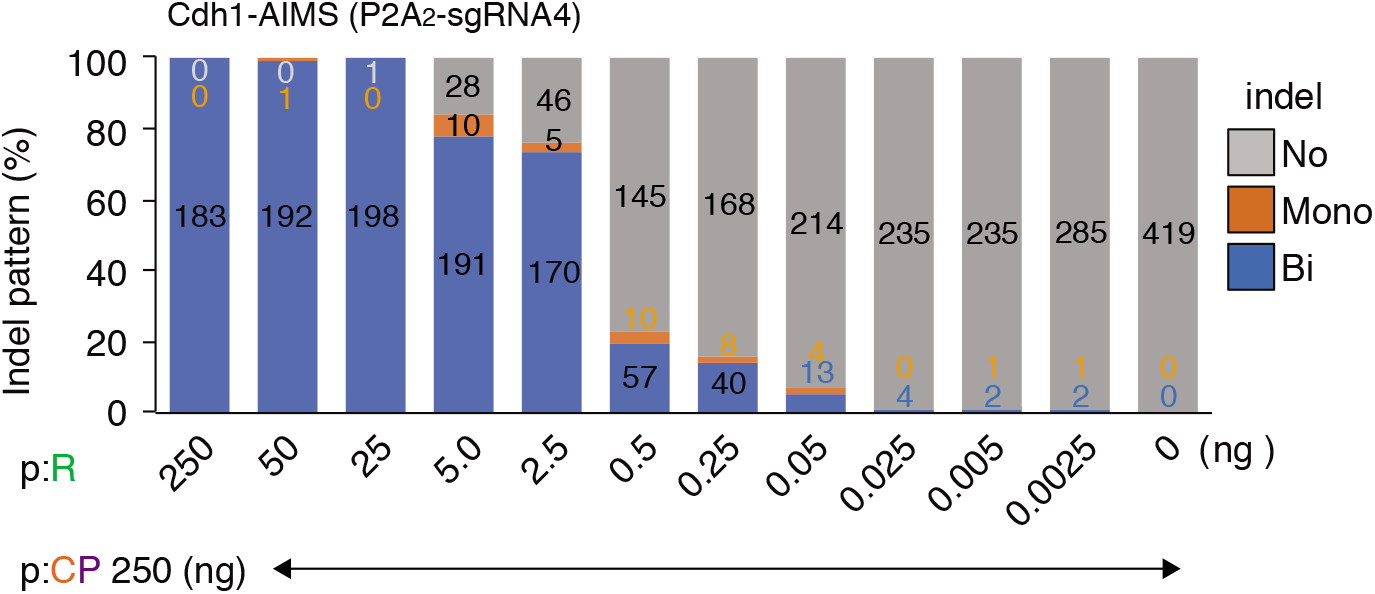
A minor effect of plasmid reduction on mono-allelic indel induction. Indel pattern is analyzed using Cdh1-P2A2-AIMS. The spacerless-PX459 plasmid (p:CP, 250 ng) is co-transfected with different amounts of the P2A2-sgRNA4 sgRNA expression plasmid (p:R). Data are shown as mean ± s.e.m. from *n* = 3 independent experiments performed at different times and total colony number analyzed is shown in each column.

**Extended Data Fig. 3.**
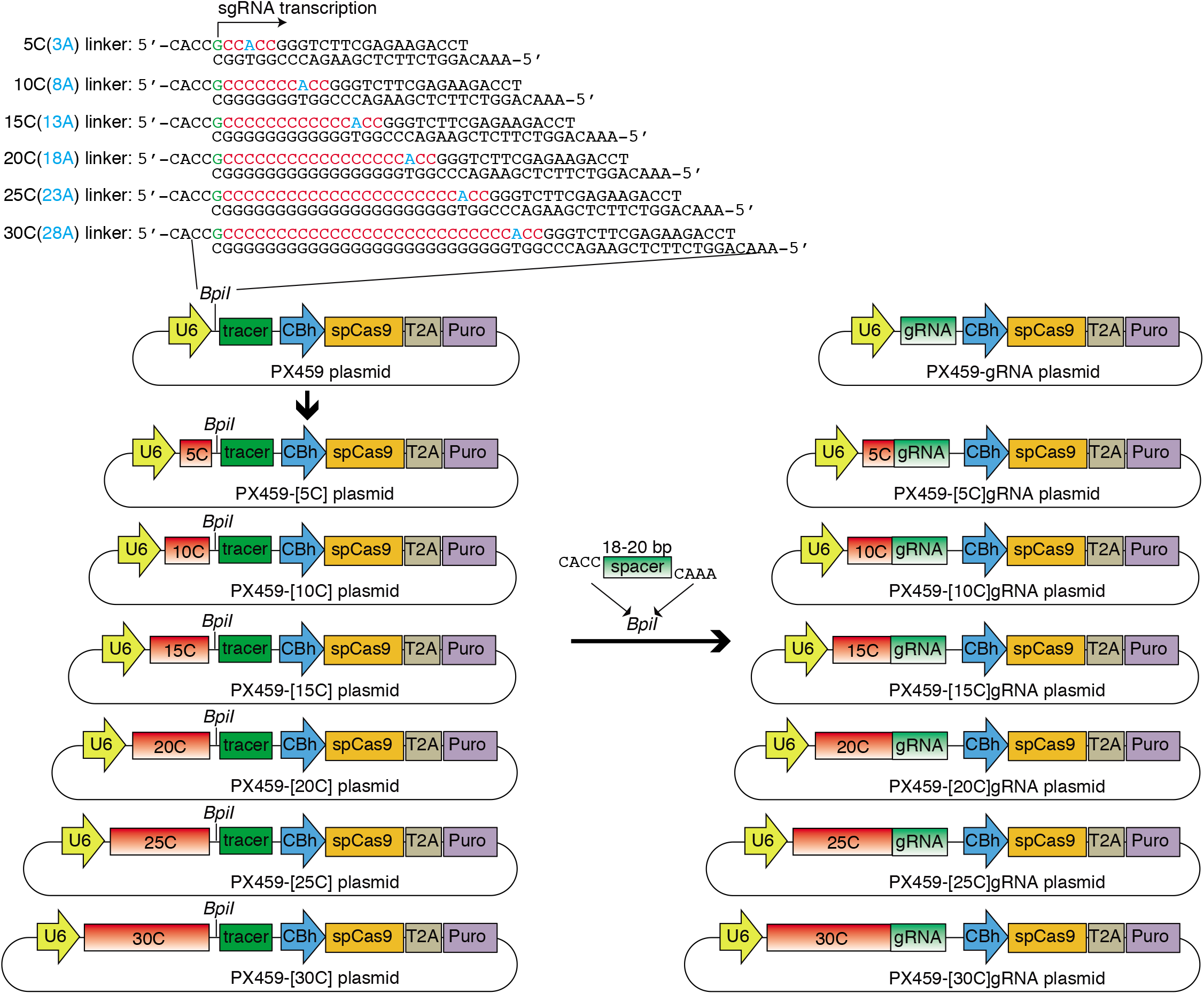
Construction of [0C]-[30C]sgRNA expressing all-in-one plasmids. The linkers are inserted into the *BpiI* site of the PX459 plasmid. Adenine (A, blue) is inserted at the third position from the 3’ end of the cytosine extension to create an overhang sequence for insertion of spacer sequences with CCAC overhang. The [5C]-[30C]sgRNA-expressing all-in-one plasmids can be produced by inserting a standard 18-20 bp spacer linker between two *BpiI* sites.

**Extended Data Fig. 4.**
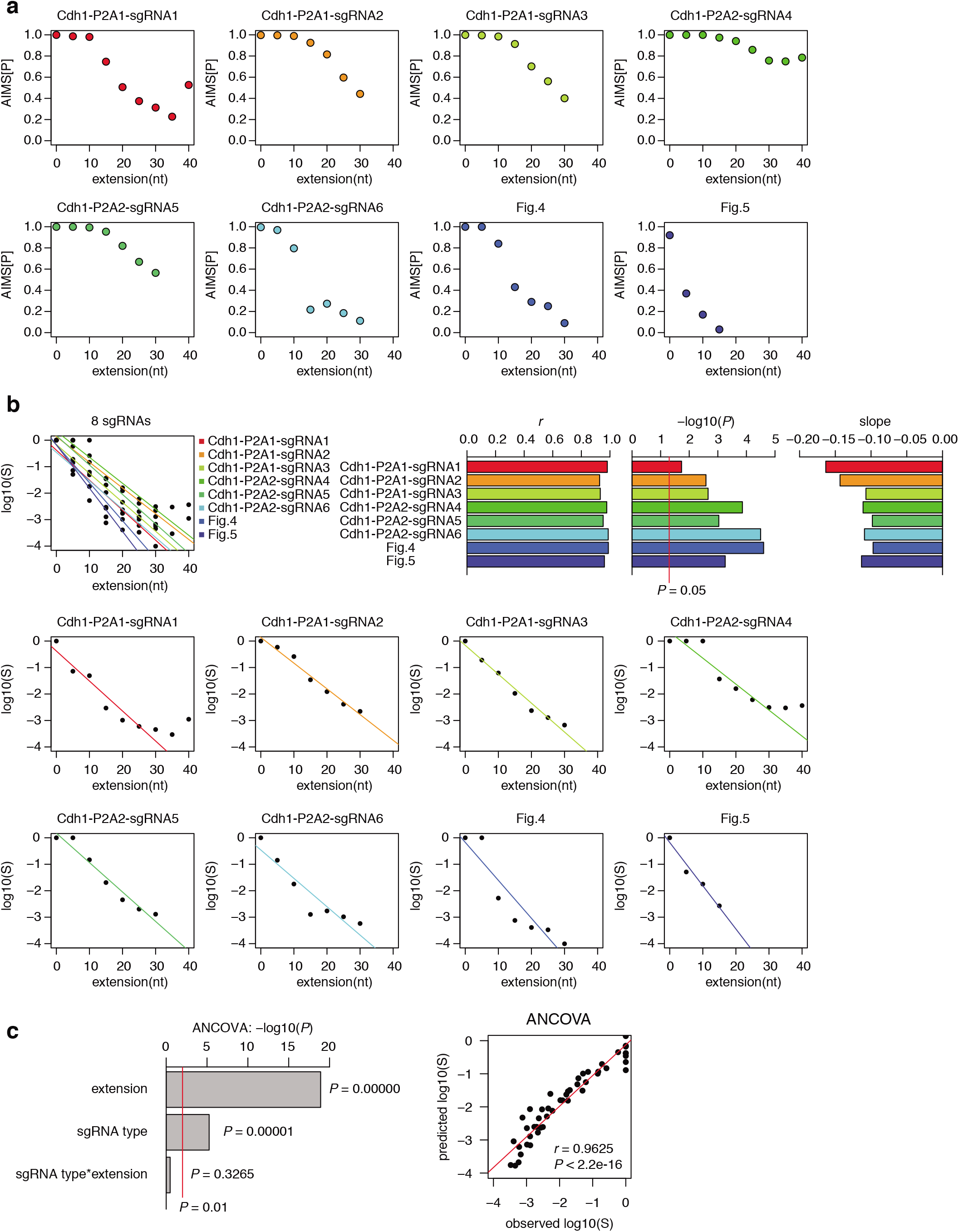
Quantitative assessment of the suppressive effects of [C] extension for different sgRNAs. **a**, Relationships between [C] extension length and AIMS[P] are shown for 8 sgRNA. **b**, Relationships between [C] extension length and concentration of effective sgRNA-Cas9 complex (log10(S)) are shown. The three upper right panels show the results of linear regression analysis, including Pearson’s correlation coefficients (*r*), P-values, and slopes. The upper left panels and 8 bottom panels show the correlation between [C] extension length and log10(S) for all sgRNAs (overlayed) and each sgRNA, respectively. Note that 8 sgRNAs have similar slope values, suggesting uniform effects of [C] extension. **c**, ANCOVA (analysis of covariance) analysis to investigate the differences of slope values for 8 sgRNAs. Statistical results for each source of variance are shown in left. Right panel shows a correlation between observed and predicted log10(S). Linear regression and Pearson’s correlation coefficient (*r*) with P-value are shown.

**Extended Data Fig. 5.**
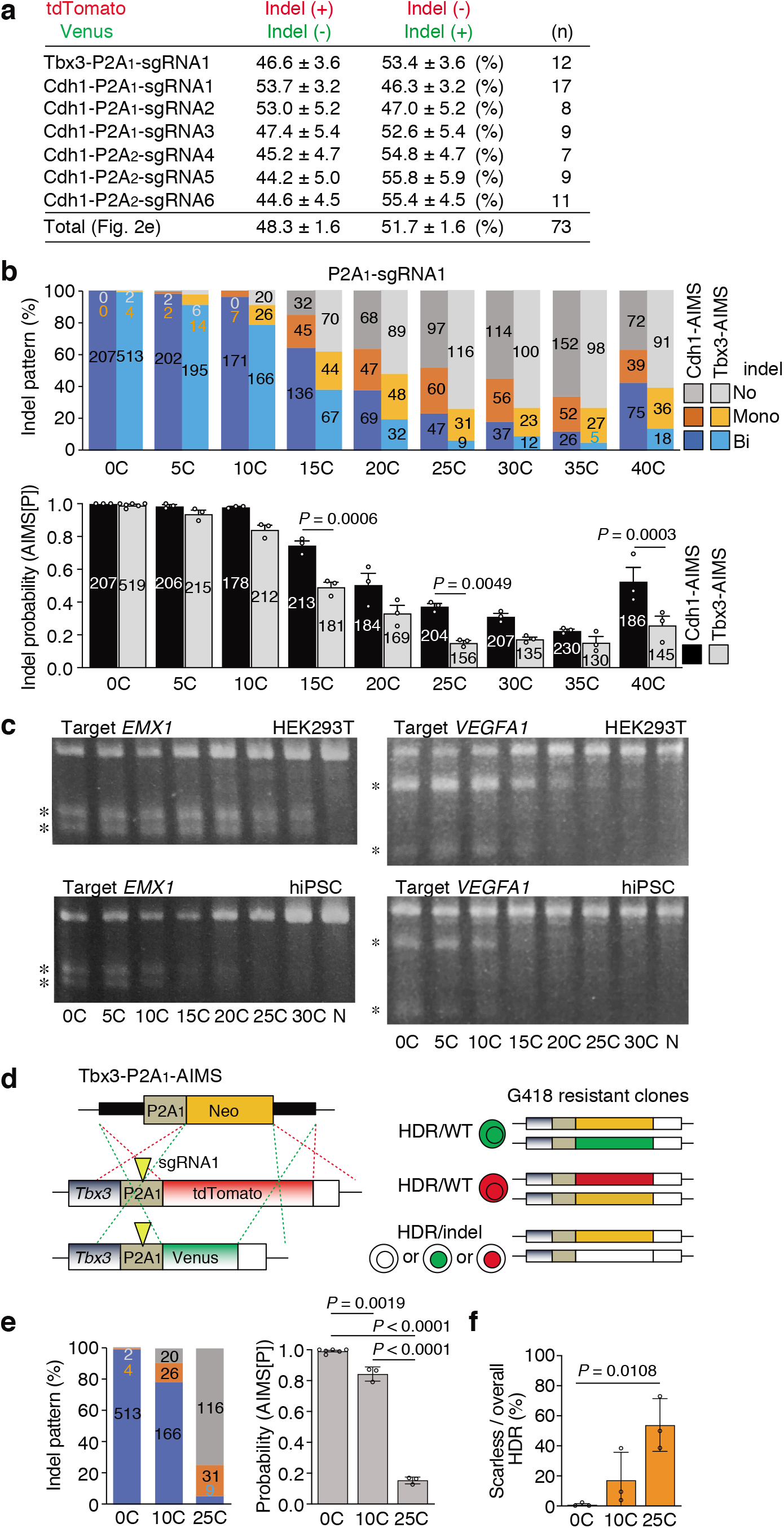
Additional results of AIMS experiments. **a**, The table shows percentages of the two types of mono-allelic indel patterns. Total indicates the mean of all data (*n* = 73), which is shown in the Fig. 2e. Data are shown as mean ± s.e.m.. **b**, Different indel frequency at different chromosomal loci. Indel pattens and probabilities (AIMS[P]) are compared between Cdh1-P2A1-AIMS and Tbx3-P2A1-AIMS. The data for the indel pattern of Cdh1-AIMS (P2A1-sgRNA1) were from Fig. 2d. Data are shown as mean ± s.e.m. from *n* = 3 (0C in Tbx3-AIMS, *n* = 6) independent experiments performed at different times and total colony number is shown in each column. Statistical significance is assessed using two-way ANOVA and post hoc Tukey–Kramer test. **c**, Comparison of indel probability between HEK293T cells and hiPSCs. Asterisks indicate PCR products digested by the T7E1 assay. N, PX459 plasmid without spacer. **d**, Schematic of measuring the frequency of scarless mono-allelic HDR without indels on the non-HDR allele using Tbx3-P2A1-AIMS. Arrowheads indicate DSB sites. **e**, Indel pattern (left) and probability (AMIS[P], right) are shown. The data of the indel pattern of Tbx3-AIMS (P2A1-sgRNA1) are from Extended Data Fig. 5b. Data are shown as mean ± s.e.m. from *n* = 3 independent experiments performed at different times and total colony number is shown in each column. Statistical significance is assessed using one-way ANOVA and post hoc Tukey–Kramer test. **f**, Frequencies of scarless HDR are shown as mean ± s.e.m. from *n* = 3 independent experiments performed at different times. Statistical significance is assessed using one-way ANOVA and post hoc Tukey–Kramer test.

**Extended Data Fig. 6.**
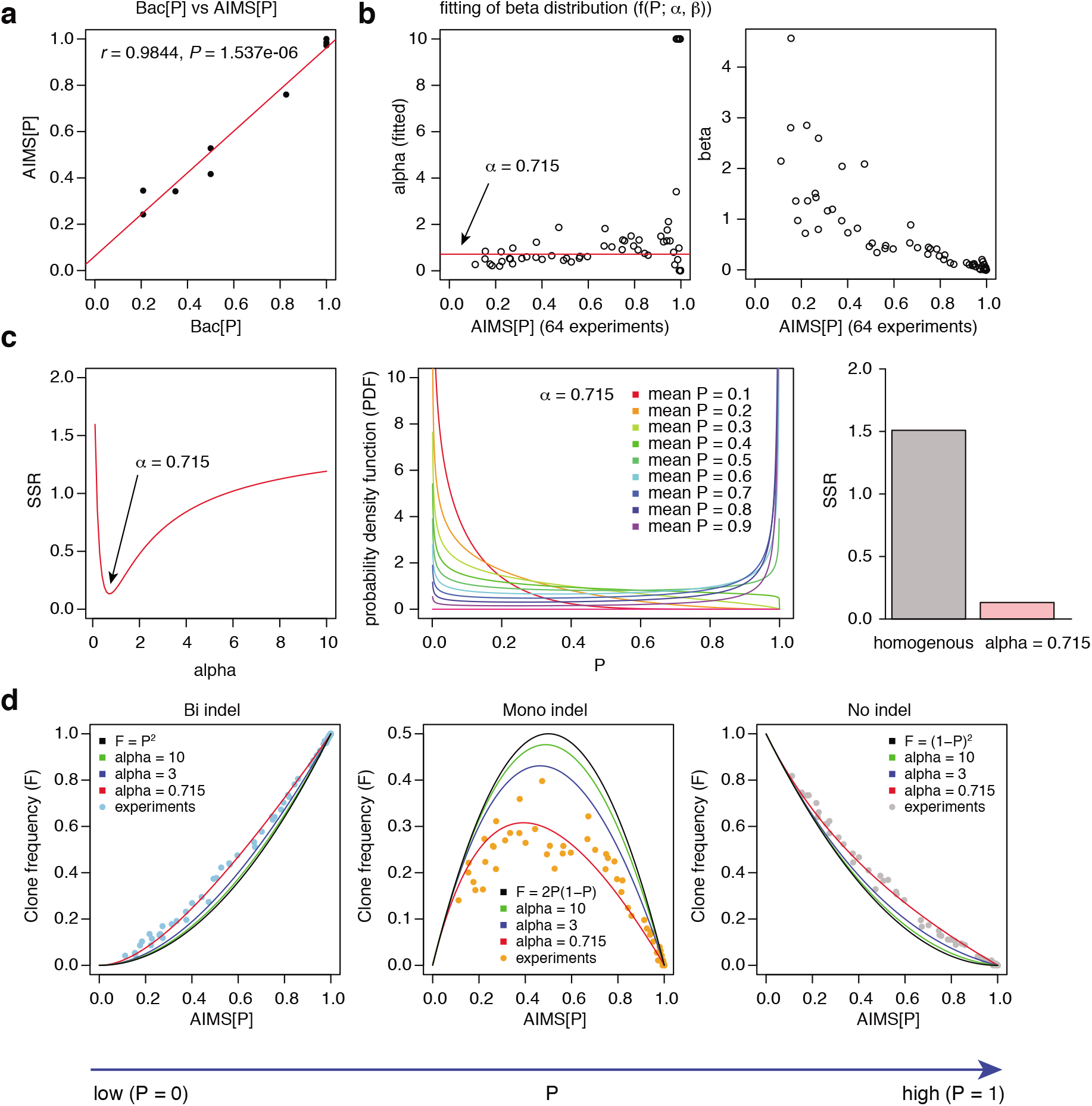
Computational modeling of single-cell heterogeneity of genome editing frequency using beta distribution. **a**, Correlation between Bac[P] and AIMS[P] shown in Fig. 3e. Linear regression and Pearson’s correlation coefficient (*r*) with P-value are shown. b, α and β values of beta distribution for each experiment that minimized sum of squared residuals (SSR) between experimental F(Bi), F(Mono), and F(No) and simulated F(Bi), F(Mono), and F(No). c, Identification of fixed α value that minimized SSR (left). Probability density functions with different mean P are shown in a middle panel. Right panel shows comparison of SSR based on the assumption that single-cell editing probability is homogenous or heterogeneous. d, Correlation between the experimental data and prediction of clone frequency for bi-, mono-, or no-indel. Black line is based on the assumption that genome editing probability is homogenous across the cell population. For the beta distribution, additional simulations in the case that α is 3 or 10 are also exhibited.

**Extended Data Fig. 7.**
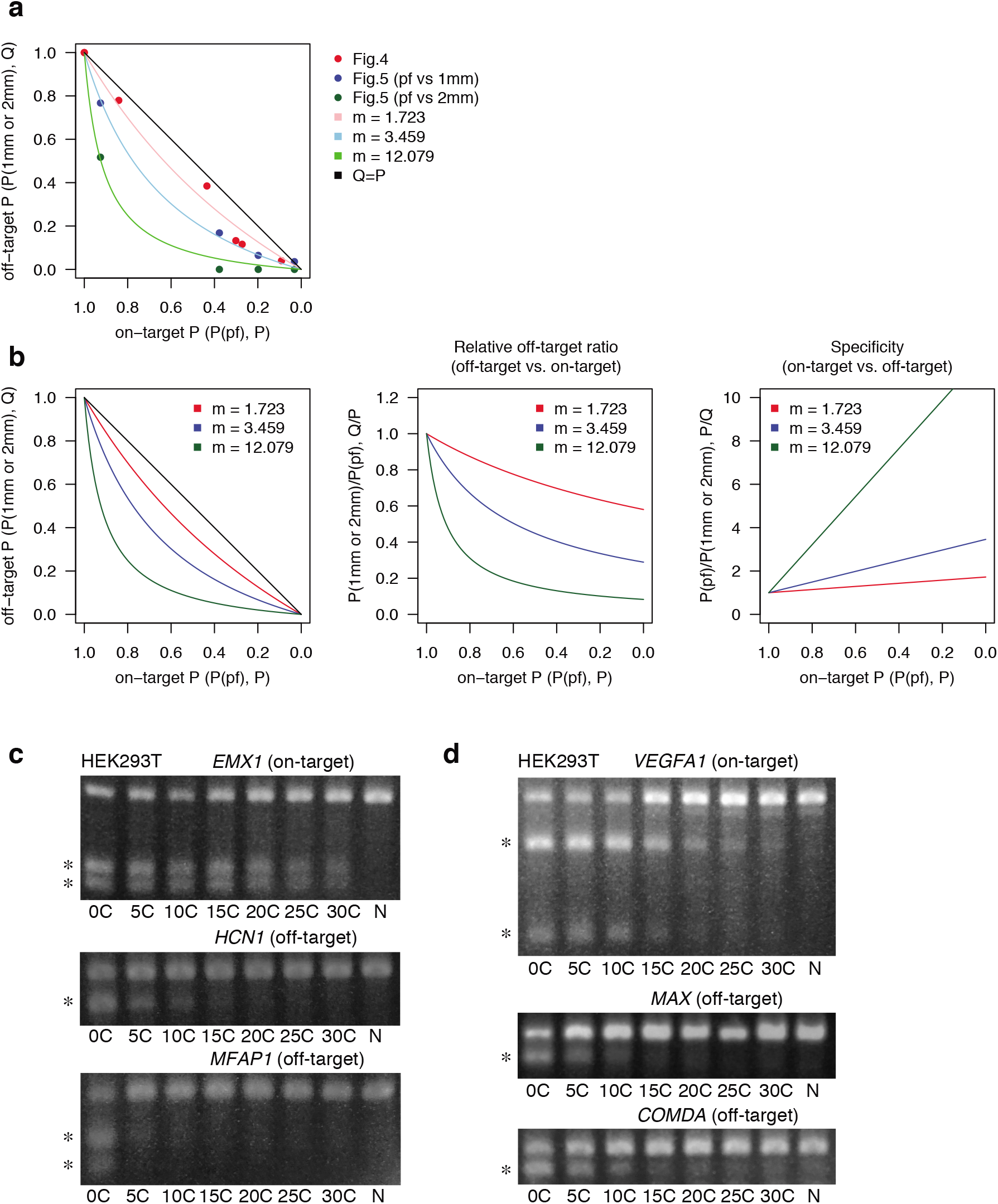
Downsizing sgRNA-Cas9 activity enhances on-target specificity and suppresses off-target effects. **a**, Relationships between on-target editing probability on perfect match target (P) and off-target probability on 1mm or 2mm target (Q) shown in Fig. 4c and Fig. 5c. Results of computational fitting are also shown. Red, blue, and green dots indicate experimental data from Fig. 4c, Fig. 5c (pf vs 1mm), and Fig. 5c (pf vs 2mm), respectively. Details of generating simulation curves are described in Methods. **b**, Computational analysis of decrease in relative off-target editing and increase in on-targeting specificity along with reduction in indel probability. pf, perfect match; 1mm, 1 bp mismatch; 2mm, 2 bp mismatch. **c, d**, Indel probability of other sgRNAs in HEK293T cells. The T7E1 assay is performed to investigate on-target and off-target indel probability for *EMX1* (**c**) and *VEGFA1* (**d**) targeting sgRNAs.

**Extended Data Fig. 8.**
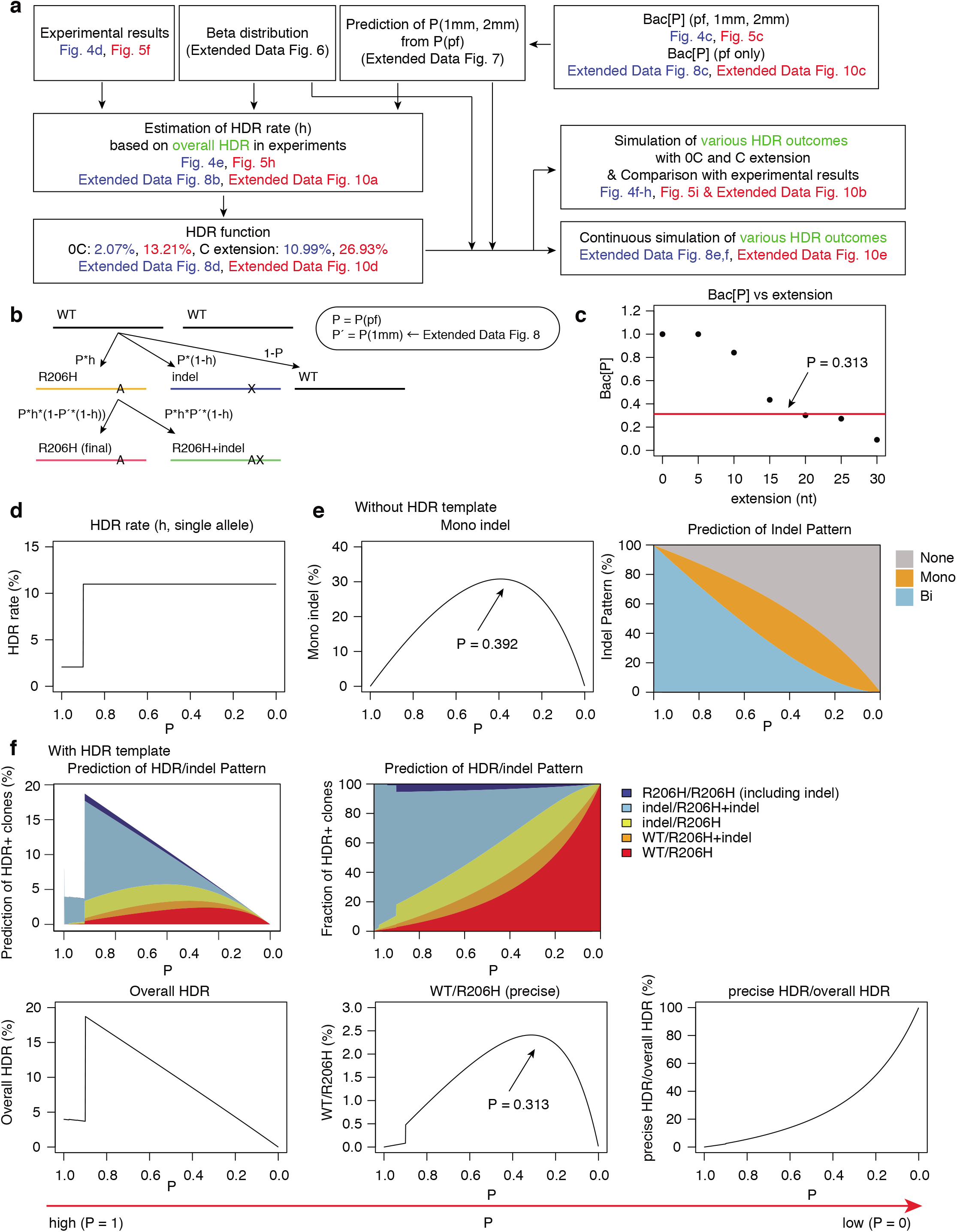
Computational simulation of precise mono-allelic HDR from homozygous states. **a**, A flow chart to establish a computational simulation of precise mono-allelic HDR to generate FOP model in mESC (Fig. 4 and Extended Data Fig. 8) or to correct a SNP in FOP hiPSCs (Fig. 5 and Extended Data Fig. 10). **b**, Scheme of HDR-mediated generation of FOP model (WT/R206H) from a WT/WT genotype. **c**, Relationship between [C] extension length and Bac[P] shown in Fig. 4c. A red line (P = 0.313) indicates the value when precise WT/R206H HDR is induced at the highest level. **d**, Hypothetical function of HDR rate (h) along indel probability (P). HDR rate is set based on the data of Fig. 4e (see details in Methods). **e**, Simulation of editing outcomes in the absence of HDR templates. Left and right panels show the relationships between indel probability (P) and frequencies for mono-allelic indel (left) or No/Mono/Bi indel (right). An arrow indicates the predicted maximum value for mono-allelic indel induction with P value of 0.392. **f**, Simulation of editing outcomes in the presence of HDR templates. Top panels show the relationships between indel probability (P) and frequencies of various HDR clones (top, left) and relative fraction (top, right). Frequencies of overall HDR and precise WT/R206H editing and the relative ratio of the WT/R206H clones (vs overall HDR) are shown in bottom panels. An arrow indicates the predicted maximum value (P = 0.313) to generate precise WT/R206H HDR clones.

**Extended Data Fig. 9.**
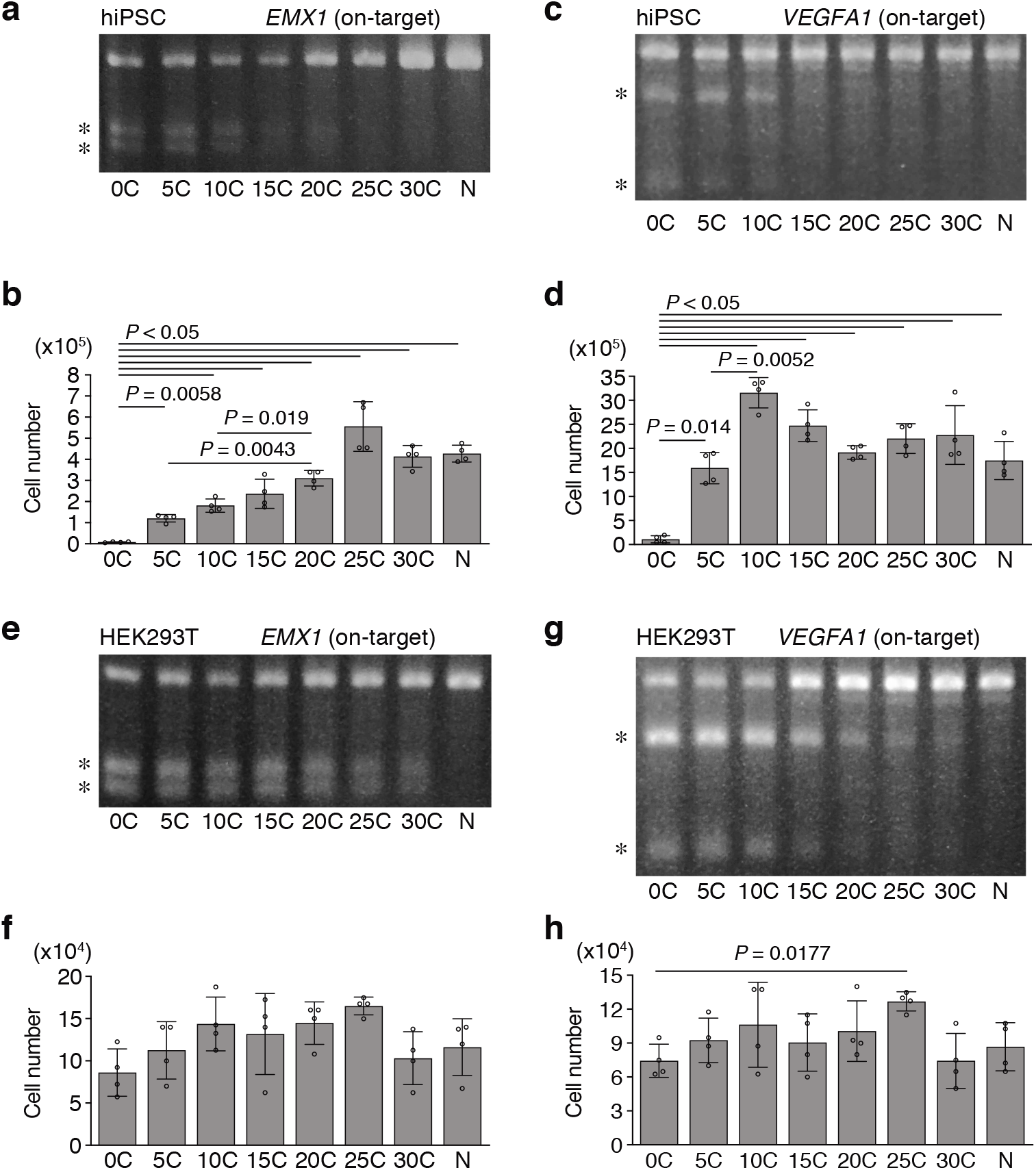
Suppression of cytotoxicity by [C] extension in hiPSCs. **a-d**, Indel probability (**a, c**) and cytotoxicity (**b, d**) of other sgRNAs are investigated in hiPSCs. The T7E1 assay is performed to investigate on-target indel probability for sgRNAs targeting *EMX1* (**a**) and *VEGFA1* (**c**). These pictures are also shown in the Extended Data Fig 5c. **e-h**, Indel probability (**e, g**) and cytotoxicity (**f, h**) are investigated in HEK293T cells. The T7E1 assay is demonstrated to investigate on-target and off-target indel probability for sgRNAs targeting *EMX1* (**e**) and *VEGFA1* (**g**). These pictures are also shown in the Extended Data Fig 7c, d. N, PX459 plasmid without spacer (**a-h**). Asterisks indicate PCR products digested by the T7E1 assay (**a, c, e, g**). Statistical significance is assessed using Welch’s test with post hoc Games–Howell test (**b, d, f, h**)

**Extended Data Fig. 10.**
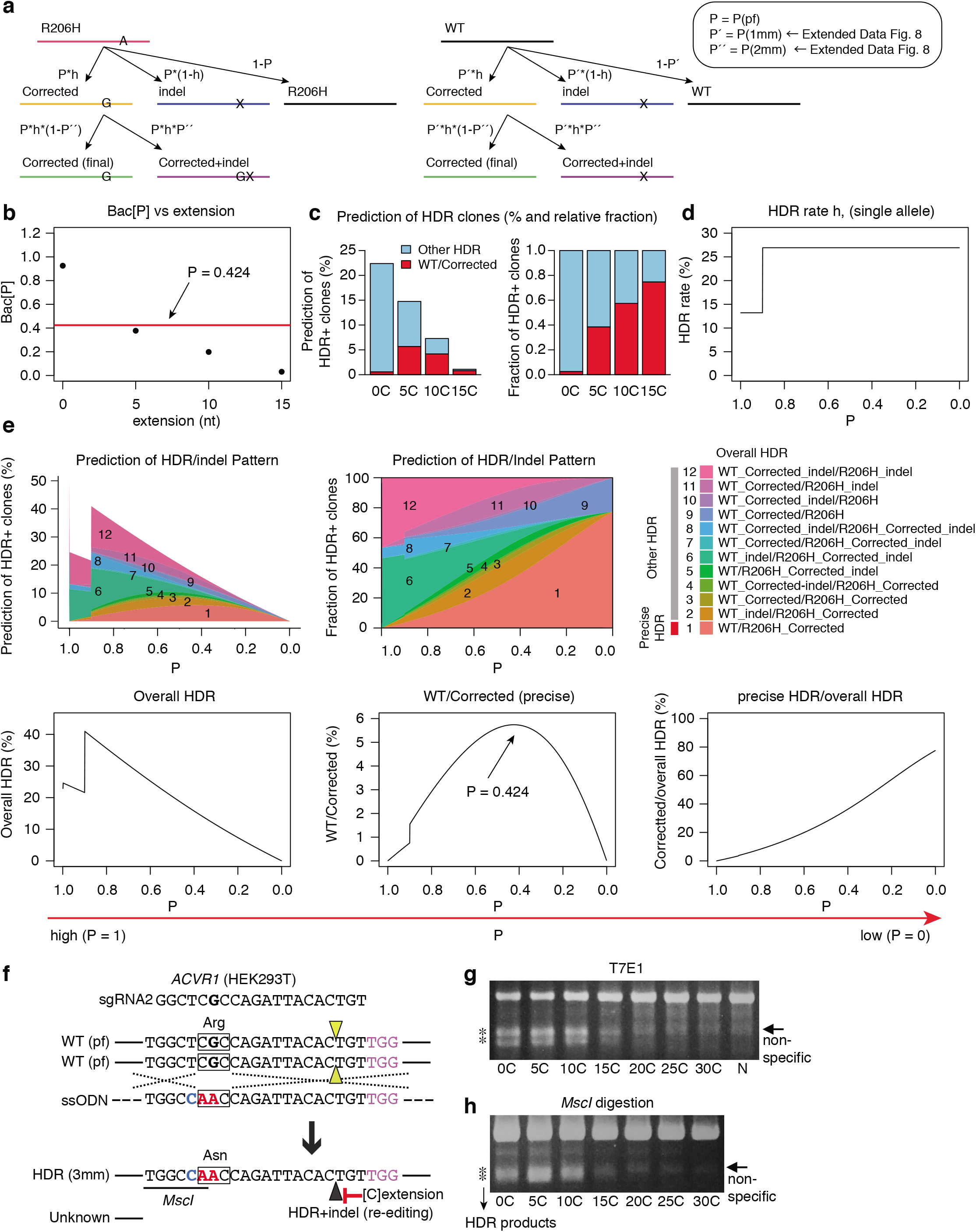
Computational simulation of precise disease gene correction. **a**, Scheme of HDR-mediated correction of FOP model (WT/R206H) to a WT/Corrected genotype. **b**, Relationship between [C] extension length and Bac[P] for R206H allele, shown in Fig. 5c. A red line (P = 0.424) indicates the value when precise WT/Corrected HDR is induced at the highest level. **c**, Prediction of precise WT/Corrected clones and other HDR clones (left) and relative fraction (right). **d**, Hypothetical function of HDR rate (h) along indel probability (P). HDR rate is set based on the data of Fig. 5h (see details in Methods). **e**, Simulation of a relationship between indel probability (P) and various HDR outcomes (top, left) and relative fraction (top, right). Frequencies of overall HDR and WT/Corrected HDR and the relative ratio of the WT/Corrected clones (vs overall HDR) are shown in bottom panels. An arrow indicates the predicted maximum value (P = 0.424) to generate precise WT/Corrected HDR clones. **f-h**, Measuring HDR frequency in HEK293T cells. Schematic of HDR for 3 bp substitution in exon 5 of *ACVR1* (**f**). Silent mutation with cytosine (C, blue) and missense mutations with two adenines (AA, red) creates a *MscI* restriction enzyme site, which allows for rapid quantification of HDR frequency. Squares indicate codon. pf, perfect match; 3mm, 3 bp mismatches. Arrowheads indicate DSB sites. Asterisks indicate PCR products digested by T7E1 (**g**) or digested by *MscI* restriction enzyme (**h**). N, PX459 plasmid without spacer.

## Main References

1 Jinek, M. et al. A programmable dual-RNA-guided DNA endonuclease in adaptive bacterial immunity. Science 337, 816–821, doi:10.1126/science.1225829 (2012).

2 Cong, L. et al. Multiplex genome engineering using CRISPR/Cas systems. Science 339, 819–823, doi:10.1126/science.1231143 (2013).

3 Mali, P. et al. RNA-guided human genome engineering via Cas9. Science 339, 823–826, doi:10.1126/science.1232033 (2013).

4 Cho, S. W., Kim, S., Kim, J. M. & Kim, J. S. Targeted genome engineering in human cells with the Cas9 RNA-guided endonuclease. Nat Biotechnol 31, 230–232, doi:10.1038/nbt.2507 (2013).

5 Haapaniemi, E., Botla, S., Persson, J., Schmierer, B. & Taipale, J. CRISPR-Cas9 genome editing induces a p53-mediated DNA damage response. Nat Med 24, 927–930, doi:10.1038/s41591-018-0049-z (2018).

6 Ihry, R. J. et al. p53 inhibits CRISPR-Cas9 engineering in human pluripotent stem cells. Nat Med 24, 939–946, doi:10.1038/s41591-018-0050-6 (2018).

7 Kosicki, M., Tomberg, K. & Bradley, A. Repair of double-strand breaks induced by CRISPR-Cas9 leads to large deletions and complex rearrangements. Nat Biotechnol 36, 765–771, doi:10.1038/nbt.4192 (2018).

8 Soldner, F. et al. Parkinson-associated risk variant in distal enhancer of alpha-synuclein modulates target gene expression. Nature 533, 95–99, doi:10.1038/nature17939 (2016).

9 Paquet, D. et al. Efficient introduction of specific homozygous and heterozygous mutations using CRISPR/Cas9. Nature 533, 125–129, doi:10.1038/nature17664 (2016).

10 Komor, A. C., Kim, Y. B., Packer, M. S., Zuris, J. A. & Liu, D. R. Programmable editing of a target base in genomic DNA without double-stranded DNA cleavage. Nature 533, 420–424, doi:10.1038/nature17946 (2016).

11 Nishida, K. et al. Targeted nucleotide editing using hybrid prokaryotic and vertebrate adaptive immune systems. Science 353, doi:10.1126/science.aaf8729 (2016).

12 Gaudelli, N. M. et al. Programmable base editing of A*T to G*C in genomic DNA without DNA cleavage. Nature 551, 464–471, doi:10.1038/nature24644 (2017).

13 Anzalone, A. V. et al. Search-and-replace genome editing without double-strand breaks or donor DNA. Nature 576, 149–157, doi:10.1038/s41586-019-1711-4 (2019).

14 Lee, H. K. et al. Targeting fidelity of adenine and cytosine base editors in mouse embryos. Nat Commun 9, 4804, doi:10.1038/s41467-018-07322-7 (2018).

15 Kim, H. S., Jeong, Y. K., Hur, J. K., Kim, J. S. & Bae, S. Adenine base editors catalyze cytosine conversions in human cells. Nat Biotechnol 37, 1145–1148, doi:10.1038/s41587-019-0254-4 (2019).

16 Lin, Q. et al. Prime genome editing in rice and wheat. Nat Biotechnol 38, 582–585, doi:10.1038/s41587-020-0455-x (2020).

17 Liu, Y. et al. Efficient generation of mouse models with the prime editing system. Cell Discov 6, 27, doi:10.1038/s41421-020-0165-z (2020).

18 Kim, J. H. et al. High cleavage efficiency of a 2A peptide derived from porcine teschovirus-1 in human cell lines, zebrafish and mice. PLoS One 6, e18556, doi:10.1371/journal.pone.0018556 (2011).

19 Russell, R. et al. A Dynamic Role of TBX3 in the Pluripotency Circuitry. Stem Cell Reports 5, 1155–1170, doi:10.1016/j.stemcr.2015.11.003 (2015).

20 Pieters, T. et al. p120 Catenin-Mediated Stabilization of E-Cadherin Is Essential for Primitive Endoderm Specification. PLoS Genet 12, e1006243, doi:10.1371/journal.pgen.1006243 (2016).

21 Chari, R., Mali, P., Moosburner, M. & Church, G. M. Unraveling CRISPR-Cas9 genome engineering parameters via a library-on-library approach. Nat Methods 12, 823–826, doi:10.1038/nmeth.3473 (2015).

22 Maurissen, T. L. & Woltjen, K. Synergistic gene editing in human iPS cells via cell cycle and DNA repair modulation. Nat Commun 11, 2876, doi:10.1038/s41467-020-16643-5 (2020).

23 Cho, S. W. et al. Analysis of off-target effects of CRISPR/Cas-derived RNA-guided endonucleases and nickases. Genome Res 24, 132–141, doi:10.1101/gr.162339.113 (2014).

24 Mullally, G. et al. 5’ modifications to CRISPR-Cas9 gRNA can change the dynamics and size of R-loops and inhibit DNA cleavage. Nucleic Acids Res 48, 6811–6823, doi:10.1093/nar/gkaa477 (2020).

25 Arimbasseri, A. G., Rijal, K. & Maraia, R. J. Transcription termination by the eukaryotic RNA polymerase III. Biochim Biophys Acta 1829, 318–330, doi:10.1016/j.bbagrm.2012.10.006 (2013).

26 Shore, E. M. et al. A recurrent mutation in the BMP type I receptor ACVR1 causes inherited and sporadic fibrodysplasia ossificans progressiva. Nat Genet 38, 525–527, doi:10.1038/ng1783 (2006).

27 Chakkalakal, S. A. et al. An Acvr1 R206H knock-in mouse has fibrodysplasia ossificans progressiva. J Bone Miner Res 27, 1746–1756, doi:10.1002/jbmr.1637 (2012).

28 Matsumoto, Y. et al. Induced pluripotent stem cells from patients with human fibrodysplasia ossificans progressiva show increased mineralization and cartilage formation. Orphanet J Rare Dis 8, 190, doi:10.1186/1750-1172-8-190 (2013).

29 Hino, K. et al. Neofunction of ACVR1 in fibrodysplasia ossificans progressiva. Proc Natl Acad Sci U S A 112, 15438–15443, doi:10.1073/pnas.1510540112 (2015).

30 Pawluk, A. et al. Naturally Occurring Off-Switches for CRISPR-Cas9. Cell 167, 1829–1838 e1829, doi:10.1016/j.cell.2016.11.017 (2016).

31 Harrington, L. B. et al. A Broad-Spectrum Inhibitor of CRISPR-Cas9. Cell 170, 1224–1233 e1215, doi:10.1016/j.cell.2017.07.037 (2017).

32 Rauch, B. J. et al. Inhibition of CRISPR-Cas9 with Bacteriophage Proteins. Cell 168, 150–158 e110, doi:10.1016/j.cell.2016.12.009 (2017).

33 Bubeck, F. et al. Engineered anti-CRISPR proteins for optogenetic control of CRISPR-Cas9. Nat Methods 15, 924–927, doi:10.1038/s41592-018-0178-9 (2018).

34 Hynes, A. P. et al. Widespread anti-CRISPR proteins in virulent bacteriophages inhibit a range of Cas9 proteins. Nat Commun 9, 2919, doi:10.1038/s41467-018-05092-w (2018).

35 Jiang, F. et al. Temperature-Responsive Competitive Inhibition of CRISPR-Cas9. Mol Cell 73, 601–610 e605, doi:10.1016/j.molcel.2018.11.016 (2019).

36 Maji, B. et al. A High-Throughput Platform to Identify Small-Molecule Inhibitors of CRISPR-Cas9. Cell 177, 1067–1079 e1019, doi:10.1016/j.cell.2019.04.009 (2019).

37 Singh, D. et al. Mechanisms of improved specificity of engineered Cas9s revealed by single-molecule FRET analysis. Nat Struct Mol Biol 25, 347–354, doi:10.1038/s41594-018-0051-7 (2018).

38 Okafor, I. C. et al. Single molecule analysis of effects of non-canonical guide RNAs and specificity-enhancing mutations on Cas9-induced DNA unwinding. Nucleic Acids Res 47, 11880–11888, doi:10.1093/nar/gkz1058 (2019).

## Methods References

39 Yagi, M. et al. Derivation of ground-state female ES cells maintaining gamete-derived DNA methylation. Nature 548, 224–227, doi:10.1038/nature23286 (2017).

40 Okita, K. et al. A more efficient method to generate integration-free human iPS cells. Nat Methods 8, 409–412, doi:10.1038/nmeth.1591 (2011).

41 Srinivas, S. et al. Cre reporter strains produced by targeted insertion of EYFP and ECFP into the ROSA26 locus. BMC Dev Biol 1, 4, doi:10.1186/1471-213x-1-4 (2001).

